# Longitudinal Network Re-organization Across Learning and Development

**DOI:** 10.1101/2020.10.23.353094

**Authors:** Ethan M. McCormick, Sabine Peters, Eveline A. Crone, Eva H. Telzer

## Abstract

While it is well understood that the brain experiences changes across short-term experience/learning and long-term development, it is unclear how these two mechanisms interact to produce developmental outcomes. Here we test an interactive model of learning and development where certain learning-related changes are constrained by developmental changes in the brain against an alternative development-as-practice model where outcomes are determined primarily by the accumulation of experience regardless of age. Participants (8-29 years) participated in a three-wave, accelerated longitudinal study during which they completed a feedback learning task during an fMRI scan. Adopting a novel longitudinal modeling approach, we probed the unique and moderated effects of learning, experience, and development simultaneously on behavioral performance and network modularity during the task. We found nonlinear patterns of development for both behavior and brain, and that greater experience supported increased learning and network modularity relative to naïve subjects. We also found changing brain-behavior relationships across adolescent development, where heightened network modularity predicted improved learning, but only following the transition from adolescence to young adulthood. These results present compelling support for an interactive view of experience and development, where changes in the brain impact behavior in context-specific fashion based on developmental goals.

## 1. Introduction

The brain is a dynamic system capable of reshaping itself across time to adapt to its external environment. For some developmental processes (e.g., cognitive control or risk-taking; Casey, 2015; or socioemotional development; Blakemore & Mills, 2014), these changes unfold across long time horizons (e.g., months or years). However, functional development does not require years, or even months, to show measurable changes. Indeed, a broad literature has demonstrated that brain function rapidly adapts to task demands and feedback to support skill acquisition or goal-directed behavior (e.g., Daw et al., 2006; Bassett et al., 2011; McCormick & Telzer, 2017a; 2017b; 2018; Telesford et al., 2017; Gerraty et al., 2018). However, it remains unclear to what extent these short-term, learning-related changes in brain activation overlap with the long-term, maturational plasticity seen across years and decades of development (Galván, 2010). Here, we test two potential explanations for how experience and development interact across time to explain changes in learning performance and the functional brain systems that support that performance across time. To probe these interactions, we adopt a novel application of longitudinal modeling that allows us to consider changes across minutes, years, and the course of development simultaneously. This approach offers an integrated perspective of learning and development as co-dependent processes of neural and behavior plasticity which interact across time.

While traditionally thought of as a period of vulnerability (Steinberg et al., 2008; Casey et al., 2008; Shulman et al., 2016), adolescence is also a period associated with increases in flexible behavior and the capacity to learn from feedback in the environment (Johnson & Wilbrecht, 2011; Crone & Dahl, 2012; Casey, 2015; Vigilant et al., 2015), with neural changes associated with age supporting increased learning (Van Duijvenvoorde et al., 2008; Peters et al., 2016; McCormick & Telzer, 2017a; Peters & Crone, 2017). In general. the ability to learn and engage in other complex cognitive tasks (Casey et al., 2005; Luna et al., 2010), improves with age through the first decades of life. However, this co-occurrence does not by itself imply that maturation is necessary for the age-related improvements in learning seen during development. With increased age also comes more experience and practice at skills needed to support task performance. Under this view, development involves the accumulation of practice or training of neural systems, and the neural mechanisms for this process should closely resemble those involved in short-term learning. In concrete terms, this would imply that developmentally younger individuals can be trained to perform as well as older individuals given sufficient practice.

In contrast, an interactive view of learning and development would suggest that certain kinds of neural changes in response to learning are constrained by developmental changes in the brain. In other words, it should be practically impossible to train a child to perform at adult levels because they have not experienced the maturational changes in the brain necessary to support that performance. This would suggest that certain kinds of neural changes in response to learning will be relatively unique to older individuals. These two alternative accounts are not mutually exclusive, since training studies in younger individuals clearly demonstrate that there is some capacity to improve cognitive performance and shift brain function even in the maturing brain (Jolles & Crone, 2012). However, the first explanation of how learning and development interact would predict that this capacity to train should be quite extensive, whereas the second explanation would predict that biological maturation imposes stricter limitations on the ability to train young individual to “adult” levels of performance. It is important to note, however, that these limitations may not be maladaptive, but rather serve some other developmental function where “immature” brain states or behavioral performance are important for flexible learning and adaptation (e.g., Johnson & Wilbrecht, 2011; Jolles & Crone, 2012; Crone & Steinbeis, 2017; McCormick & Telzer, 2017a).

A major challenge in modeling experience and development simultaneously is that in real data, they are often confounded (Bell, 1953; Jolles & Crone, 2012; Telzer et al., 2018) in developmental models. In longitudinal studies which use a cohort-sequential (or panel) design, where individuals are repeatedly assessed at the same ages, older participants are also more-experienced participants (both in life and practice in the specific measures of interest). In neuroimaging contexts, these experience effects can confound developmental effects in a number of ways, including reducing anxiety about the scanner environment, changing baseline conditions (the “task B” problem), or reduce errors on tasks through familiarity rather than change in underlying ability (Jolles & Crone, 2012; Telzer et al., 2018). Fortunately, we can leverage an alternative, the accelerated longitudinal design, to address these challenges. In accelerated longitudinal studies, individuals vary in the age of first assessment and are followed longitudinally thereafter. By adopting this design, we can de-couple experience from age (or other measure of developmental stage) sufficiently to successfully model the accumulation of experience and developmental maturation simultaneously (McCormick, *preprint*).

The current study tests the two competing hypotheses of how experience and development interact to drive neural plasticity during learning. Participants across a wide age range (8-29 years) participated in a three-wave, accelerated longitudinal neuroimaging study during which they completed a feedback learning task. By leveraging the accelerated longitudinal design and a novel extension of mixed-effects models (McCormick, *preprint*), we differentiate between three temporal levels of neural plasticity: 1) short-term practice-related changes within a scan session (within-individual) across blocks of feedback learning; 2) long-term changes within individuals, across measurement occasions (i.e., waves); and 3) the mixed (i.e., within- and between-individual) effect of changes associated with age. By considering these three levels in the same model, we can partition effects at each level. (1) Within-session changes reflect how brain and behavior adapt during learning the task structure, (2) between-session changes reflect changes due to experience after repeated exposure to the task and testing environments, (3) while age reflects the developmental effect. Importantly, including effects at the second level allows us to de-confound age and experience, giving a more reliable estimate of the developmental effect. Because learning is an integrative process, involving the interactions between many brain regions (Bassett et al., 2011; Gerraty et al., 2014; Bassett et al., 2015; McCormick & Telzer, 2017a; Gerraty et al., 2018; McCormick, Gates, & Telzer, 2019), we test this developmental model in the context of brain networks. Specifically, we model the interaction of practice, experience, and development effects on network modularity. Modularity is a measure of the degree of network segregation into distinct functional units (Bullmore & Bassett, 2011). Higher levels of modularity in brain networks predicts increased learning (Bassett et al., 2011; Ellefsen et al., 2015) and working memory (Braun et al., 2015) performance in adults.

Our analytic approach to addressing these questions involved several steps. First, we fit mixed-effects models with only linear and quadratic effects of age on behavioral performance and network modularity during learning separately (Peters et al., 2016; Peters & Crone, 2017) for comparison to more complex models. We then included predictors of within- and between-session change as main effects to consider the unique effects of practice, experience, and development, before fitting a model that included interaction terms between our predictors. This third model allowed us to probe how the effects of the lower-level predictors change across development in a continuous fashion. Finally, we estimated a brain-as-predictor model where we probed how brain states (e.g., high versus low modularity) differentially predicted learning performance across practice, experience, and development. This final model tests a core difference between the two explanations of developmental improvement in learning performance. In the development-as-practice view, network modularity should predict learning performance consistently regardless of when in the developmental trajectory (i.e., there is no moderation by age). This is contrasted by the interaction view of development and experience, where we would expect that modularity would predict performance differentially depending on age.

## 2. Methods

### 2.1 Sample

A total of 299 participants (ages 8-29 years; 153 female) participated in a 3-wave, accelerated longitudinal MRI study. Participants were scanned every 2 years, spanning a 5-year period (Figure 1). At wave 1, 28 participants were excluded for a number of factors including not completing the MRI session (N=4), excessive movement during the scan session (>3 mm relative motion in any direction/rotation) (N=22), ADD diagnosis disclosure (N=1), and reported medicine use (N=1), resulting in a final sample of 271 participants at the initial data collection (140 female; *M*_*age*_=14.17, *SD*=3.63, *range*=8.01–25.95 years). At wave 2 (2 years later), 254 participants were scanned (33 could not be scanned due to braces; 11 declined to return). Of the scanned participants, an additional 21 were excluded (12 for motion; 2 for preprocessing errors; 5 for T2 artifacts; 1 for medicine use; 1 for ADD diagnosis), leaving a final sample of 233 participants (121 female; *M*_*age*_=16.15, *SD*=3.62, *range*=10.02–26.61 years). During the final wave (2 years later), 243 participants were scanned (11 could not be scanned due to braces; 45 declined to return). Of these, 11 were excluded (3 did not complete MRI session; 4 for motion; 2 for processing errors; 1 for medicine use; 1 for ADD diagnosis), for a total final sample of 232 participants (121 female; *M*_*age*_=18.15, *SD*=3.68, *range*=11.94–28.72 years). Across the dataset, 183 participants had data at all three waves, 78 participants had data at two waves, 31 participants had data at only one wave, and 7 were excluded at all three waves. A total of 736 scans were included for final data analyses. When considered at the trail level, these scans yielded 4799 observations for modeling.

**Figure 1.**
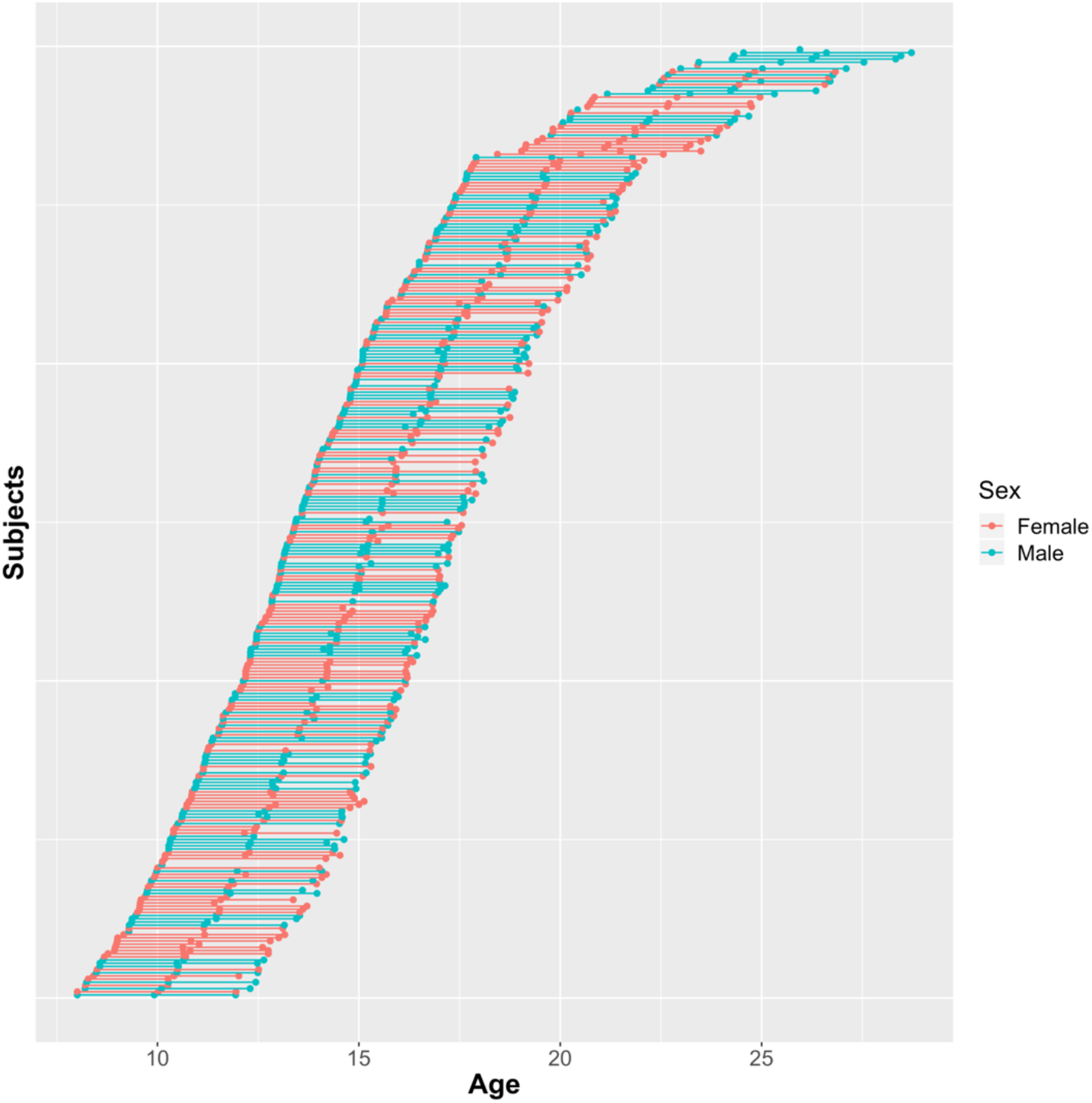
Structure of repeated measures within the accelerated longitudinal design. Participants are ordered in ascending order based on their age at wave 1. Sex is denoted by separate colors.

IQ scores were measured at the first two waves of data collection, using the WISC-III (for participants < 16 years; *N*_*W1*_ = 195; *N*_*W2*_ = 119) or WAIS-III (for participants ≥ 16 years; *N*_*W1*_ = 76; *N*_*W2*_ = 114). All participants were within the normal range at wave 1 (*M*=109.8, *SD*=10.34, *range*=80-142.5) and wave 2 (*M*=108.3, *SD*=10.27, *range*=80-147.5). Further details and the distributions of the descriptive variables are available in the supplemental material.

### 2.2 Feedback Learning Task

Participants completed a feedback learning task during an fMRI session (Peters et al., 2014; Peters et al., 2016). On each trial, participants saw a screen with three empty boxes and one (out of a possible set of three) stimulus underneath (Figure 2). Participants were told that each stimulus within a given set had a corresponding correct location among the empty boxes and that their goal on the task was to appropriately sort each stimulus into its location. For each stimulus-location choice, participants either received positive (a “+” sign) or negative (a “−” sign) feedback based on their choice. Positive feedback indicated correct stimulus placement, while negative feedback indicated incorrect placement. Each stimulus within a set associated with a unique, deterministically correct location. Stimuli within a set were presented in pseudorandom order, constrained such that no stimulus within a set was present more than twice in a row. After a maximum of 12 trials per block, or after all three stimuli within a set were correctly placed twice (indicating that all locations were successfully learned), stimulus sets were swapped out for a new set with three new stimuli. Participants saw a total of 15 blocks of 3-stimuli sets (for a maximum possible of 180 trials) at waves 1 and 2, and 10 blocks (maximum possible 120 trials) at wave 3. Prior to the MRI session, participants practiced three example sets of stimuli. Each trial consisted of the following: 1) a 500-ms fixation cross, 2) stimulus presentation for 2500 ms while participants made location decisions, and 3) feedback presentation for 1000 ms. Trials were separated by intervals jittered based on OptSeq (Dale, 1999), with durations that varied between 0 and 6 s.

**Figure 2.**
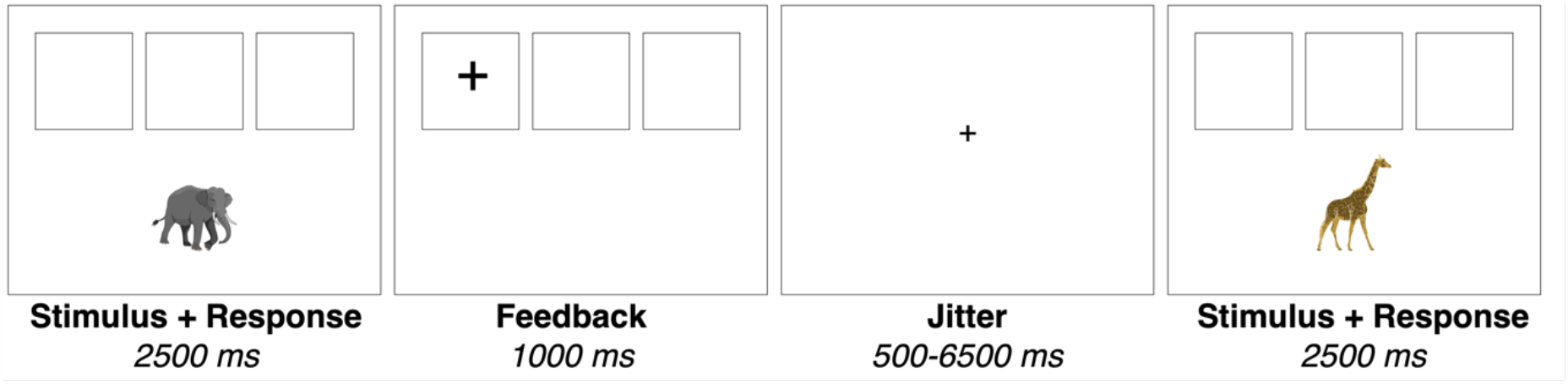
During the Feedback Learning task, participants learned the correct placement of each stimuli (e.g., the elephant) through feedback. Participants received either positive or negative feedback on each trial.

### 2.3 Behavioral Analyses

#### 2.3.1 Task Metrics of Behavior

Our primary metric of task performance was the learning rate participants displayed in forming correct stimulus-location associations. To calculate learning rate, we distinguished between two phases of task performance: the learning and the application phase (Peters et al., 2014; Peters et al., 2016). The learning phase was defined as trials where the correct location for a given stimulus was still unknown, and participants needed to rely on trial-and-error or hypothesis testing to correctly place the stimulus. Trials in the learning phase could result in either positive (indicating a future stay strategy) or negative (prompting a future shift strategy) feedback. In contrast, the application phase was defined as trials where the correct location for the presented stimulus is already known (as established by an earlier learning trial) and participants correctly place that stimulus again. Learning rate was calculated as the proportion of trials in the learning phase where feedback was correctly applied in the following trial involving the same stimulus (either as repeated placement following positive feedback or as altered placement following negative feedback) out of all the trials during the learning phase.

#### 2.3.2 Linear Mixed-Effects Model

To test our developmental/experience interaction model, we fit a linear mixed-effects model to participants’ learning rate data. We followed a model-building procedure similar to the one used in previous work in this sample (Peters et al., 2016; Peters & Crone, 2017). This procedure involved a build-up approach where we tested main effects and then interactions of time to establish the optimal developmental form before bringing in additional predictors. We fit a random effects ANOVA model with a random intercept which served as a comparison for subsequent models. For descriptive purposes at the random effects level, we fit a three level model where blocks were nested within wave and then within person, however for comparison with future models, we also fit a two level model where wave and age were included at level 1. To compare across levels of change, we constructed a model using the lme4 software package through R (version 1.1-21; Bates et al., 2015), where stimulus blocks (N=1-max 15) and wave (N=1-3) were nested within individual, and age was included as a time-varying covariate. Because wave (i.e., repeated exposure to the task) was a predictor of interest, we did not nest with respect to wave since that would result in a variable that acts as both a nesting factor and linear effect of interest. We included interactions between wave, age, and blocks. To capture more complex changes in behavior between blocks of the task, we utilized piece-wise regression at level 1 (Flora, 2008; Li et al., 2009), including predictors which model the linear effects across the first and second half of the task separately. This model resulted in the following equation:

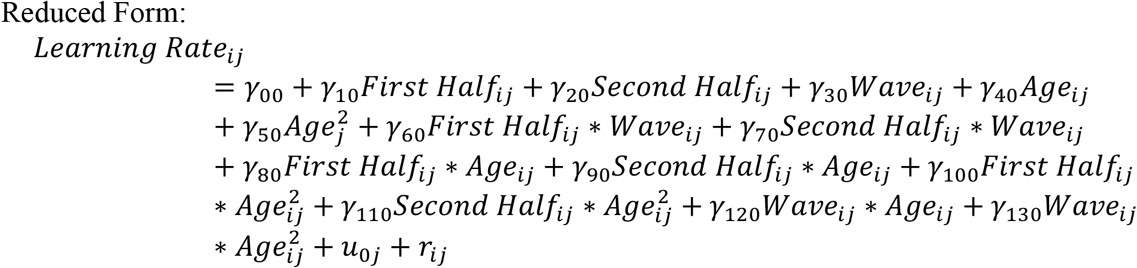

While previous work has discouraged using wave as a predictor in longitudinal models (instead using precise age; see Mehta & West, 2000), here we draw a meaningful distinction between wave and age. We would expect changes in behavior after each subsequent exposure the task environment (Telzer, et al., 2018), which due to the sampling method in accelerated longitudinal designs is disassociated with age to some degree because a large age range is represented at each wave.

### 2.4 fMRI Data Acquisition and Processing

#### 2.4.1 MRI data acquisition

Scans across all three waves were acquired using the same Philips 3T MRI scanner, utilizing identical scan settings. The Feedback Learning Task included T2*-weighted echoplanar images (EPI; slice thickness=2.75mm; 38 slices; sequential acquisition; TR=2.2sec; TE=30ms; FOV=220 × 220 × 114.68mm). Additionally, structural images were acquired, including a high-resolution 3D T1-FFE anatomical scan (TR=9.76ms; TE=4.59ms; 140 slices; voxel size=0.875 × 0.875 × 1.2mm; FOV=224 × 177 × 168mm; flip angle=8). Prior to undergoing the scan procedure, participants were introduced to the scanner environment (e.g., space and noises) through a mock scan session.

#### 2.4.2 fMRI data preprocessing and analysis

Preprocessing and analyses utilized a suite of tools from FSL FMRIBs Software Library (FSL v6.0; https://fsl.fmrib.ox.ac.uk/fsl/), Steps taken during preprocessing included skull stripping of all images using BET; and slice-to-slice motion correction of EPI images using MCFLIRT; co-registeration in a two-step sequence to the high-resolution T2-weighted and T1-FFE anatomical images using FLIRT in order to warp them into the standard stereotactic space defined by the Montreal Neurological Institute (MNI) and the International Consortium for Brain Mapping; and the application of a 128s high-pass temporal filter to remove low frequency drift within the time-series.

#### 2.4.3 Nuisance Regressors

Prior to modeling the fMRI data further, we took several steps to reduce the influence of motion. Motion, as measured by framewise displacement (Power et al., 2012), was minimal across the sample (mean across participants = 0.12 mm FD; max = 0.77 mm; average percentage of volumes with > 0.3 mm FD = 4.95%). We also controlled for 8 nuisance regressors in the GLM and time-series analyses: 6 motion parameters generated during realignment and the average signal from both the white matter and cerebrospinal fluid masks. Previous work (see Ciric et al., 2017) has shown that these strategies reduce the influence of motion on functional connectivity analyses.

#### 2.4.4 Graph Construction

We then utilized a graph theoretical approach to investigate how networks in the brain changed across levels of practice, experience, and development. Using a subset of the BigBrain parcellation scheme (Sietzman et al., 2020), an atlas comprised of 300, 5-mm sphere parcels from cortical and subcortical regions, we extracted functional timeseries data for each block (15 in total) in order to model changes in network structure across time during the task. We chose to examine network features between regions with theoretical relevance to task performance during learning. This resulted in 147 regions including those in the cingulo-opercular (14), default mode (55), fronto-parietal (27), salience (14), ventral (9) and dorsal (14) attention, hippocampal (6), and reward (8) sub-networks (for the relevant ROI coordinates on a whole-brain projection, see Figure S1). Selection of these networks were guided by those regions engaged in the feedback learning task in previous research (Peters et al., 2016; Peters & Crone, 2016) or classically engaged during learning and decision-making (Daw & Shohamy, 2008; Sadaghiani & D’Esposito, 2014; McCormick & Telzer, 2018a; McCormick et al., 2019). This subset was chosen to balance including enough regions of interest with the challenges of computing whole-brain networks on relatively short timeseries. Regions were included or excluded as a group based on their network label (e.g., all fronto-paretial regions were included while all visual regions were excluded).

To extract, we constructed a task regressor made from the onset and duration of each block of stimuli convolved with an HRF function. These regressors were multiplied with the entire timeseries extracted from ROIs in order to give a set of time-series files for each individual at each wave. Block durations (*M* = 45.32 s; *SD* = 3.26; *range* = 37.27-60.23 s) were comparable to approaches used in dynamic functional connectivity analyses (approximately 30s; Shirer et al., 2012; Gonzalez-Castillo et al., 2015). Correlation matrices were constructed by computing the zero-lag cross-correlation between each ROI. Graph metrics were calculated across a range of costs (5-20% in 5% increments; Cohen & D’Esposito, 2016). We utilized the standard community assignment for distinguishing within-versus between-network edges (Sietzman et al., 2020).

#### 2.4.5 Graph Metric

For our measure of brain development, we calculated network modularity, or the degree to which communities within the brain network are segregated. Modularity is computed as the relative number of edges between nodes of the same community compared to the number of total edges within the brain graph. We calculated modularity (Q*) using positive and negative weighted edges as:

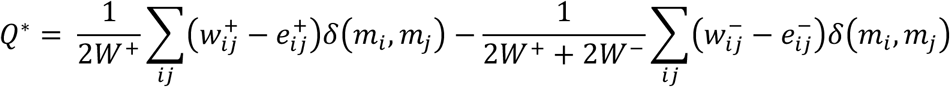

where w^+^ is the number of positively weighted edges and w^−^ is the number of negatively weighted edges. The *e*_*ij*_ term represents the expected number of edges between two nodes *i* and *j*, and the *δ*(*m*_i_, *m*_*j*_) term is 1 if the nodes *i* and *j* are in the same module and 0 if they are not in the same module. Notice that negatively weighted edges are given less influence than positively weighted edges in computing modularity (Rubinov & Sporns, 2011).

### 2.5 Developmental Model

We then utilized the same multi-level modeling approach used for the behavior to model change in brain networks across blocks, waves, and age:

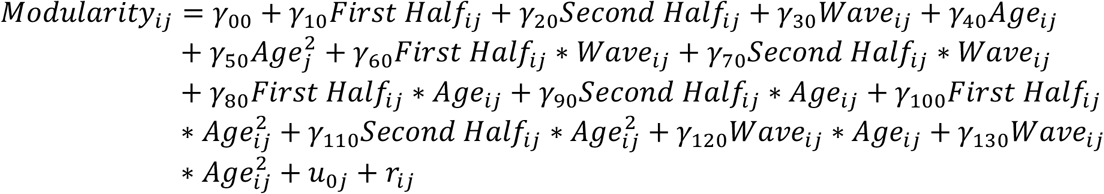

### 2.6 Probing Interactions

To better understand potential interaction effects involving age in the models of behavior and brain, we probed the effects at four distinct ages (Figure 3). Ages were chosen to be evenly spaced (approximately standard deviation distances within the sample) and roughly correspond to different developmental periods including early, middle, and late adolescences, as well as young adulthood (e.g., Shulman et al., 2016). Interaction effects in the model are continuous across the age range and therefore leverage information across the sample. However, due to lower coverage of observations at later ages, the simple slope estimates when probing the interaction at these levels have correspondingly larger standard errors.

**Figure 3.**
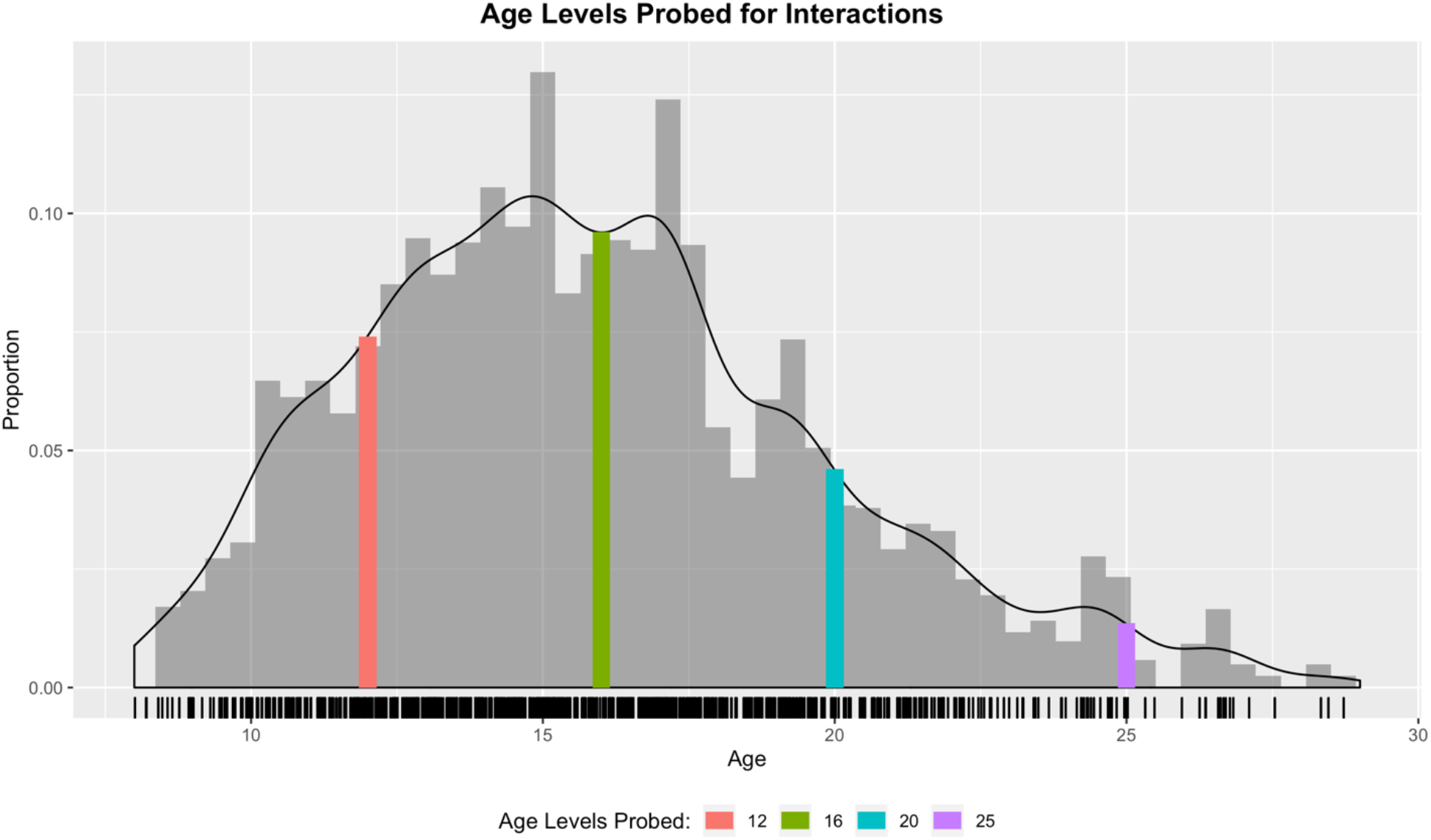
Probing Interactions with Age. Four distinct ages were chosen to probe interactions with age, including during early adolescence (12, [~ -1SD]; red), middle adolescence (16, [~ mean age], green), late adolescence (20, [+1SD]; blue), and young adulthood (25, [~ +2SD], purple). Bar height represents the proportion of observations at that level. Rug plot hashes (below x-axis) represent individual observations.

## 3. Results

### 3.1 Descriptives and Age-only Growth Models

Before formally fitting models to the data, we assessed descriptives of both learning rate and network modularity as a function of wave and age (taking the mean of within-session data). Connected data points represent the same individual across time and waves are labeled with different colors (Figure 4A & B). As reported earlier (Peters & Crone, 2017), learning rate was high overall with late adolescents performing near ceiling, and either leveling off or declining for older participants. Although not constrained in the same way as learning performance at upper values, neural network modularity appears to increase at earlier ages and declining at later ages (see Supplemental for formal regions of significance analysis for all models). As our first model building step, we fit a relatively simple model by including linear and quadratic effects of age to both learning rate and network modularity. Predicted values of each measure were obtained from the mixed-effects model (Figure 4C & D; see Table 1), confirming these trends. However, this simple model neither captures within-session practice-related change, nor does it disaggregate within-person changes due to experience (i.e., across waves) and between-person changes due to maturation (i.e., across age). We next formally tested these effects using the developmental model specified above.

**Figure 4.**
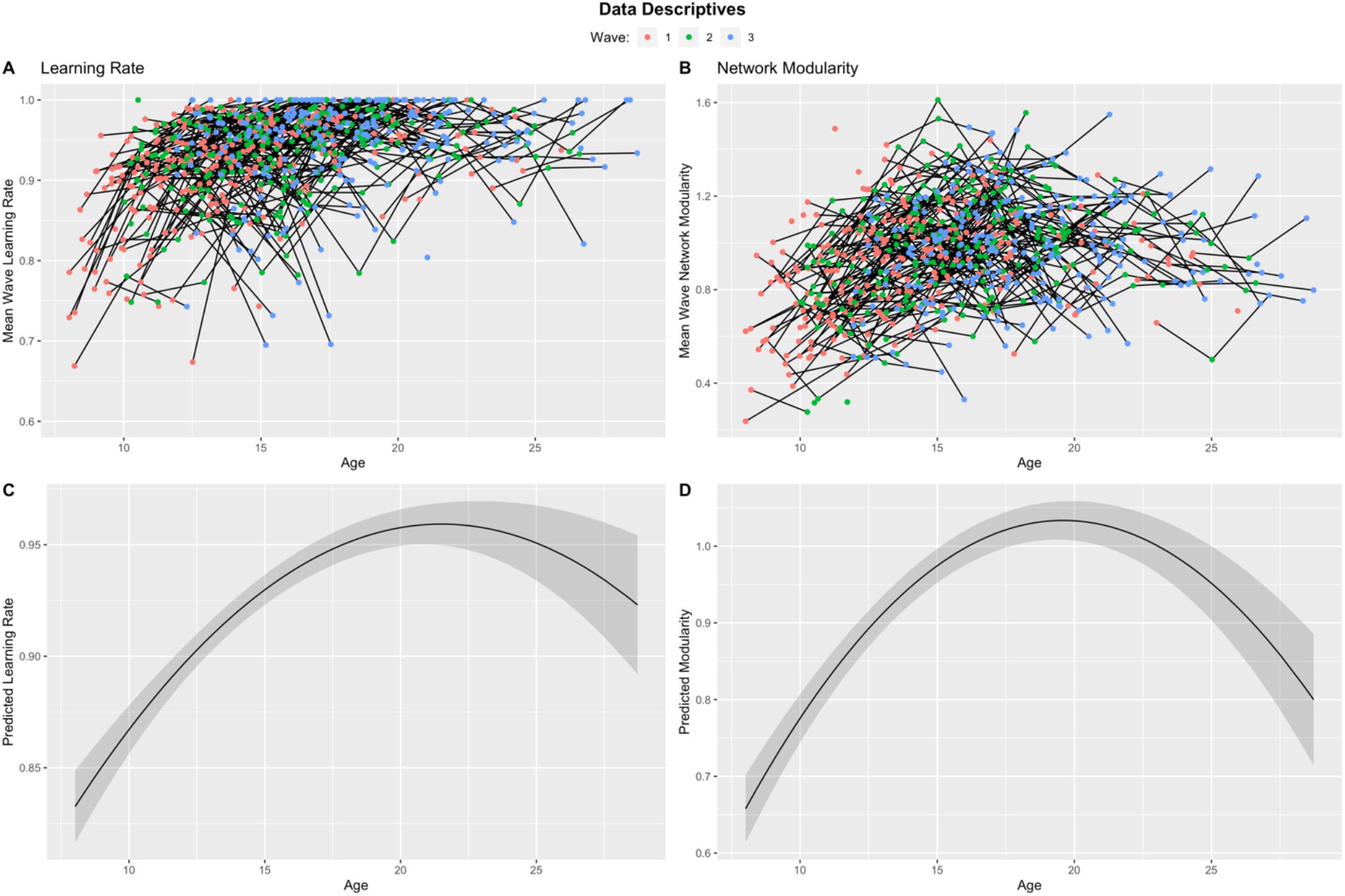
Learning (A) and Network Modularity (B) Across Age and Wave. Means of within-session data were plotted against age to visualize potential time-related trends. Wave number was indicated by color, and data from the same individual were connected by a solid line. If individuals only contributed data at one timepoint, data was indicated with a lone point. Linear mixed-effects models that only consider the mixed effect of age suggest quadratic effects peaking in late adolescence for learning performance (C), and for network modularity (D). These models fail to consider effects of experience (i.e., wave) and within-session practice effects.

**Table 1.** Model Output from Age-Only Models.

| *Predictors* | **Learning Rate** |  |  |  | **Network Modularity** |  |  |  |
| --- | --- | --- | --- | --- | --- | --- | --- | --- |
|  | *Estimates* | *SE* | *P-Value* | *df* | *Estimates* | *SE* | *P-Value* | *df* |
| Intercept | 0.937 | 0.004 | <0.001 | 359.604 | 0.994 | 0.011 | <0.001 | 361.688 |
| Age<sub>std</sub> | 0.277 | 0.023 | <0.001 | 1003.133 | 0.275 | 0.023 | <0.001 | 1992.711 |
| Age<sup>2</sup><sub>std</sub> | -0.140 | 0.021 | <0.001 | 1924.892 | -0.209 | 0.020 | <0.001 | 3523.698 |
| **Random Effects** |  |  |  |  |  |  |  |  |
| $\sigma^2$ | 0.010 | | | | 0.055 | | | |
| $\tau_{00}$ | 0.002<sub>ID</sub> | | | | 0.030<sub>ID</sub> | | | |
| ICC | 0.18 |  |  |  | 0.35 |  |  |  |
| N | 297<sub>ID</sub> |  |  |  | 297<sub>ID</sub> |  |  |  |
| Observations | 4799 |  |  |  | 4799 |  |  |  |
| Marginal R<sup>2</sup> /<br>Conditional R<sup>2</sup> | 0.065 / 0.236 |  |  |  | 0.075 / 0.402 |  |  |  |

*Note:* All effects rounded to the third decimal place for display purposes. std = standardized effects reported. $\sigma^2$ = level 1 random effect. $\tau_{00}$ = higher level random effect (effect specified by subscript). ICC = Intraclass correlation. N = number of units at each level.

### 3.2 Behavioral Improvements in Learning Performance

We began by fitting an unconditional random effects ANOVA model (i.e., a random intercept at each level) to determine the distribution of variance across the three levels of the model (level 1 = within-session; level 2 = waves; level 3 = individual). Results indicated that the majority of variance in learning rate was between trials within the same scan session (68.7%), an additional 19.3% of the variance was between scan sessions within the same individual (i.e., change across waves), and the remaining 11.9% variance was accounted for by between-individual differences in overall learning performance (see Table 2 for full details). However, to establish a baseline for future models, we also fit a two level random effects ANOVA model where level 1 and 2 are collapsed (see Table 2).

**Table 2.**
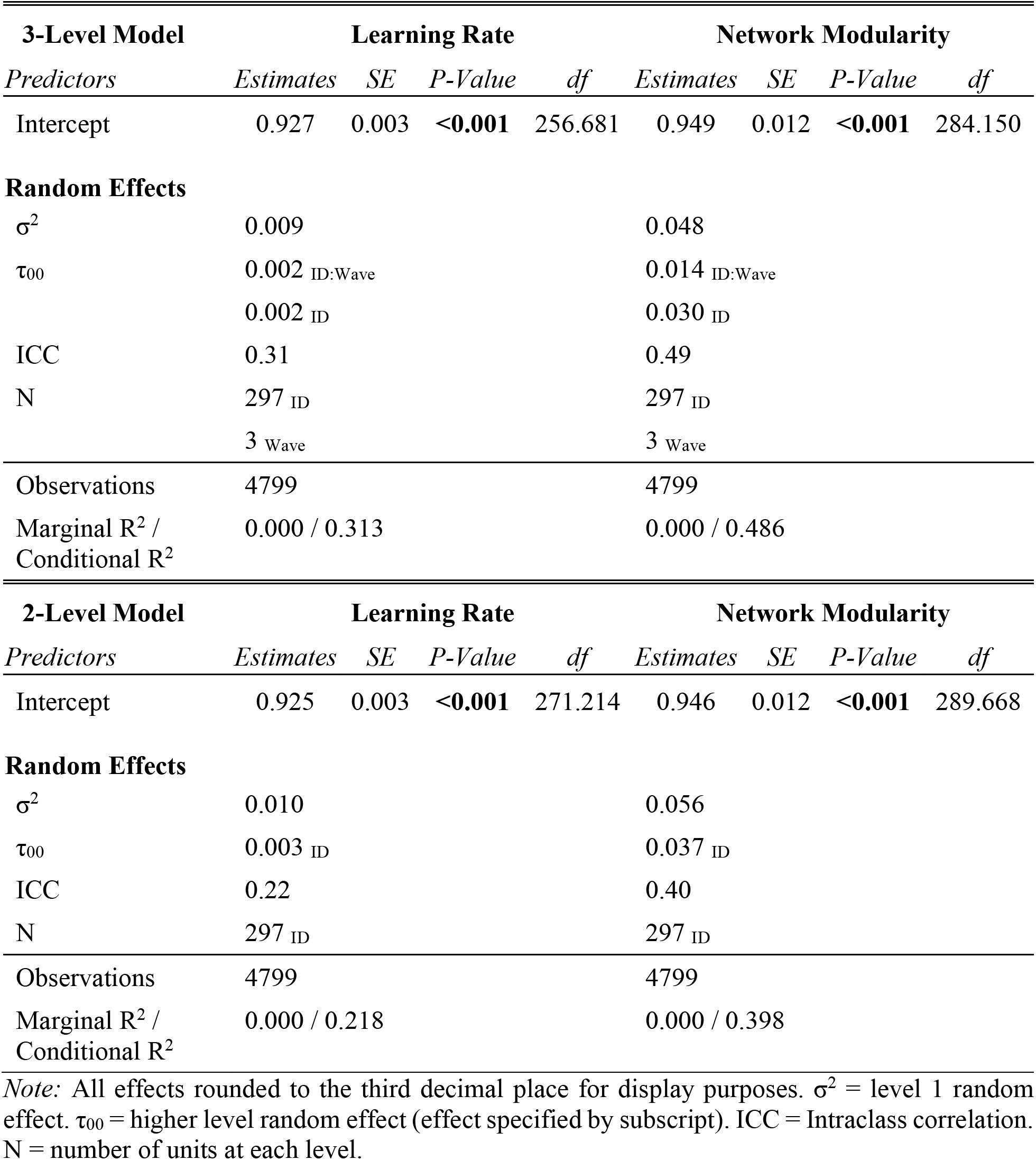
Model Output from Random Effects ANOVA Models.

#### 3.2.1 Separating Effects of Age and Wave

Next, we fit a mixed effects model with fixed predictors of learning rate at each level to assess the separable effects of age and wave on learning rate. Importantly, wave and age were sufficiently decoupled in this model (*r* = .397, *variance inflation factor* = 1.19, *SE inflation* = 1.08 times), and all predictors were centered. In this model neither effect of the within-session practice predictors were significant. This suggests that there was no total systematic change in learning rate within a scan session net the effects of wave and age, nor was there a significant within-person effect of wave. In other words, neither within-session practice or between-session experience related to increased performance when accounting for the age-related change. However, individuals showed significant linear (β = .250, *SE* = .031, *p* < .001) and quadratic (β = -.136, *SE* = .021, *p* < .001) effects of age on learning rate. All results are reported as standardized effects (Table 3). A likelihood ratio test suggests that this model offers an improvement over the unconditional model 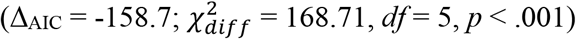.

**Table 3.** Model Output from Main Effects-Only Models.

| *Predictors* | **Learning Rate** |  |  |  | **Network Modularity** |  |  |  |
| --- | --- | --- | --- | --- | --- | --- | --- | --- |
|  | *Estimates* | *SE* | *P-Value* | *df* | *Estimates* | *SE* | *P-Value* | *df* |
| Intercept | 0.927 | 0.004 | <**0.001** | 717.508 | 0.987 | 0.013 | <**0.001** | 520.937 |
| First Half<sub>std</sub> | 0.025 | 0.016 | 0.113 | 4473.435 | 0.088 | 0.013 | <**0.001** | 4491.297 |
| Second Half<sub>std</sub> | 0.022 | 0.016 | 0.151 | 4473.468 | 0.121 | 0.013 | <**0.001** | 4491.319 |
| Wave<sub>std</sub> | 0.023 | 0.018 | 0.198 | 1205.547 | -0.004 | 0.019 | 0.845 | 694.861 |
| Age<sub>std</sub> | 0.250 | 0.31 | <**0.001** | 299.628 | 0.285 | 0.039 | <**0.001** | 311.080 |
| Age<sup>2</sup><sub>std</sub> | -0.136 | 0.021 | <**0.001** | 1941.031 | -0.209 | 0.020 | <**0.001** | 3747.423 |
| **Random Effects** |  |  |  |  |  |  |  |  |
| $\sigma^2$ | 0.010 | | | | 0.051 | | | |
| $\tau_{00}$ | 0.002<sub>ID</sub> | | | | 0.030<sub>ID</sub> | | | |
| ICC | 0.18 |  |  |  | 0.37 |  |  |  |
| N | 297<sub>ID</sub> |  |  |  | 297<sub>ID</sub> |  |  |  |
| Observations | 4799 |  |  |  | 4799 |  |  |  |
| Marginal R<sup>2</sup> /<br>Conditional R<sup>2</sup> | 0.060 / 0.232 |  |  |  | 0.112 / 0.441 |  |  |  |

*Note:* All effects rounded to the third decimal place for display purposes. std = standardized effects reported. $\sigma^2$ = level 1 random effect. $\tau_{00}$ = higher level random effect (effect specified by subscript). ICC = Intraclass correlation. N = number of units at each level.

#### 3.2.2 Learning Rate Improvements Show Interactions Across Levels of Time

Finally, we tested the interactive model of learning and development for participants’ learning rates. To do so, we added two-way cross-level interaction terms to the previous model. Three-way interactions were explored but were not found to be significant and so the model with two-way interactions was retained. All predictors were centered to create interaction terms which were uncorrelated with the main effects and to facilitate the interpretation of main effects in the presence of interaction terms (Aiken & West, 1991). There was a significant positive interaction of wave and the quadratic effect of age on learning rate (β = .083, *SE* = .021, *p* < .001) such that at each later waves, the quadratic decreases lessen. To probe this interaction, we plotted mean within-session level increases in learning rate across age for each wave (Figure 5). Results suggest that without repeated exposure to the task (i.e., experience) there are predicted decreases in learning performance at younger ages (red trajectory), but that practice helps compensate and cause performance to level off instead (green and blue trajectories). The quadratic effect where wave is coded as zero (i.e., wave 2) reflects the effects seen in the age-only model and is consistent with prior research (Peters et al., 2016; Peters & Crone, 2017), however, by including interactions with experience, we show how that total effect is influenced by repeated exposure to the task (see Table 4 for full details). A likelihood ratio test suggests that this model offers an improvement over the main-effects only model 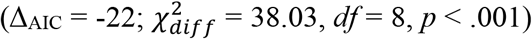. In a follow-up sensitivity analysis, we found that this pattern of effects held when including IQ as a covariate.

**Figure 5.**
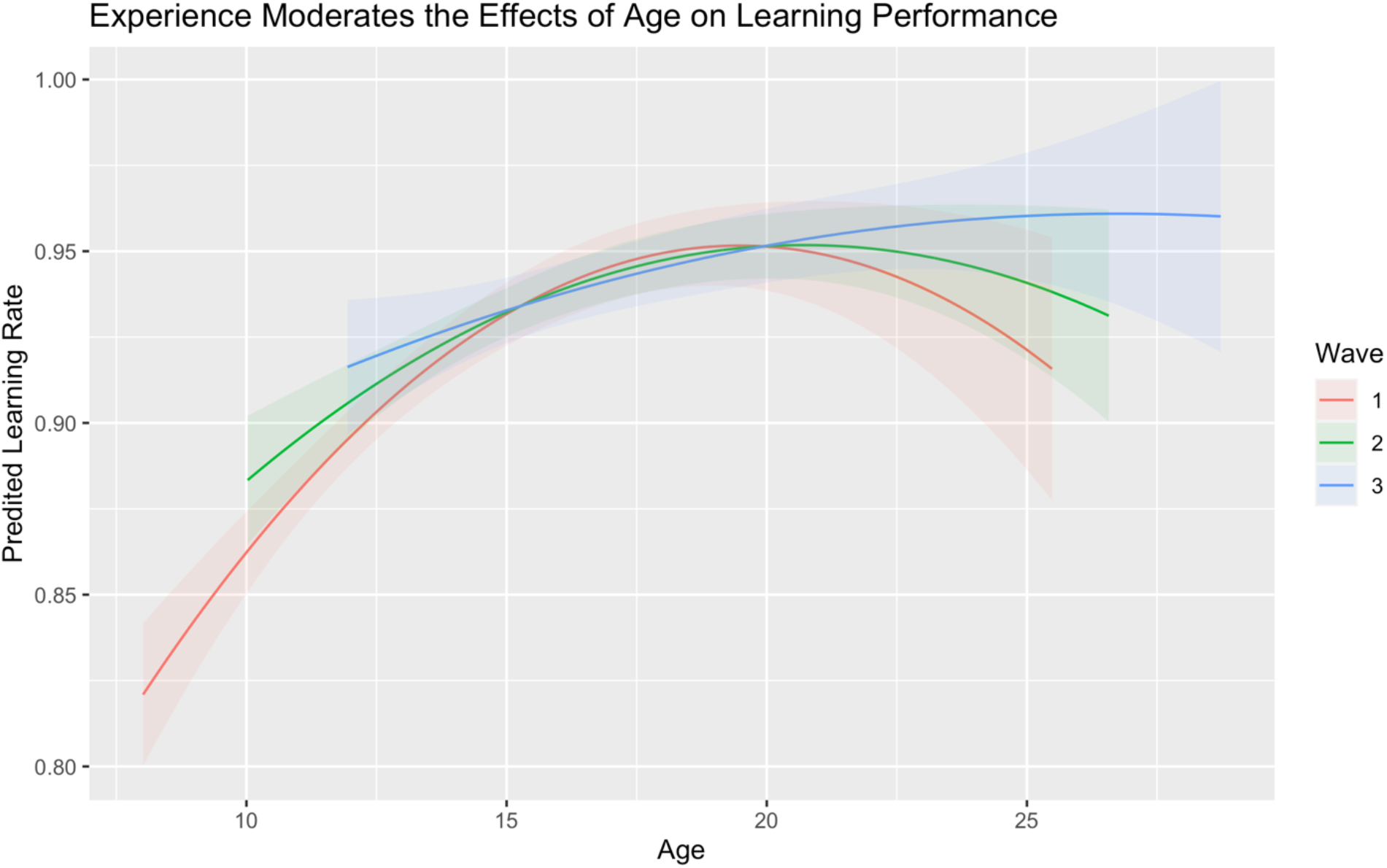
Increased Experience Impacts Learning Trajectories. Compared with first exposure (red), accumulating experience (green and blue) between waves tended to predict better learning performance at later ages, compensating for expected declines in performance during young adulthood.

**Table 4.** Model Output from Interactions Models.

| *Predictors* | **Learning Rate** |  |  |  | **Network Modularity** |  |  |  |
| --- | --- | --- | --- | --- | --- | --- | --- | --- |
|  | *Estimates* | *SE* | *P-Value* | *df* | *Estimates* | *SE* | *P-Value* | *df* |
| Intercept | 0.935 | 0.005 | <0.001 | 768.901 | 0.997 | 0.014 | <0.001 | 513.739 |
| First Half<sub>std</sub> | -0.007 | 0.020 | 0.751 | 4473.838 | 0.094 | 0.017 | <0.001 | 4495.715 |
| Second Half<sub>std</sub> | 0.021 | 0.020 | 0.297 | 4473.787 | 0.115 | 0.017 | <0.001 | 4495.776 |
| Wave<sub>std</sub> | 0.011 | 0.031 | 0.722 | 3143.488 | -0.044 | 0.029 | 0.131 | 1734.974 |
| Age<sub>std</sub> | 0.160 | 0.044 | <0.001 | 650.191 | 0.327 | 0.049 | <0.001 | 471.938 |
| Age<sup>2</sup><sub>std</sub> | -0.080 | 0.042 | 0.053 | 615.163 | -0.261 | 0.047 | <0.001 | 513.285 |
| First Half * Wave<sub>std</sub> | -0.016 | 0.022 | 0.469 | 4484.108 | -0.036 | 0.019 | 0.058 | 4508.580 |
| Second Half * Wave<sub>std</sub> | -0.044 | 0.022 | 0.048 | 4484.488 | -0.022 | 0.019 | 0.244 | 4508.591 |
| First Half * Age<sub>std</sub> | -0.049 | 0.022 | 0.027 | 4466.136 | 0.068 | 0.019 | <0.001 | 4482.736 |
| Second Half * Age<sub>std</sub> | 0.025 | 0.022 | 0.252 | 4466.143 | -0.004 | 0.019 | 0.818 | 4482.736 |
| Wave * Age<sub>std</sub> | -0.038 | 0.024 | 0.111 | 870.145 | 0.002 | 0.026 | 0.947 | 648.001 |
| First Half * Age<sup>2</sup><sub>std</sub> | 0.047 | 0.024 | 0.048 | 4464.637 | -0.027 | 0.020 | 0.182 | 4481.364 |
| Second Half * Age<sup>2</sup><sub>std</sub> | -0.025 | 0.024 | 0.292 | 4464.630 | -0.002 | 0.020 | 0.906 | 4481.360 |
| Wave * Age<sup>2</sup><sub>std</sub> | 0.083 | 0.021 | <0.001 | 4746.733 | 0.057 | 0.018 | 0.001 | 4674.686 |
| **Random Effects** |  |  |  |  |  |  |  |  |
| $\sigma^2$ | 0.010 | | | | 0.051 | | | |
| $\tau_{00}$ | 0.002<sub>ID</sub> | | | | 0.030<sub>ID</sub> | | | |
| ICC | 0.19 |  |  |  | 0.37 |  |  |  |
| N | 297<sub>ID</sub> |  |  |  | 297<sub>ID</sub> |  |  |  |
| Observations | 4799 |  |  |  | 4799 |  |  |  |
| Marginal R<sup>2</sup> /<br>Conditional R<sup>2</sup> | 0.066 / 0.239 |  |  |  | 0.126 / 0.449 |  |  |  |

*Note:* All effects rounded to the third decimal place for display purposes. std = standardized effects reported. $\sigma^2$ = level 1 random effect. $\tau_{00}$ = higher level random effect (effect specified by subscript). ICC = Intraclass correlation. N = number of units at each level.

### 3.3 Changes in Network Modularity

Similar to the behavioral analysis, we first fit an unconditional random effects ANOVA model to participants’ neural network modularity data. The majority of variance in network modularity was between trials within the same scan session (51.4%), relatively less (15.6%) of the variance was between scan sessions within the same individual (i.e., change across waves), with the remainder (33.0%) accounted for by between-individual differences in network modularity (Table 2).

#### 3.3.1 Separating Effects of Age and Wave

Next, we fit a main-effects only model with predictors including task block, wave, and age. There were linear (β = .285, *SE* = .0.39, *p* < .001) and quadratic (β = -.136, *SE* = .021, *p* < .001) effects of age, such that network modularity tended to increase early in adolescence and level off and decrease across late adolescence and young adulthood. Additionally, modularity increased across blocks within waves (i.e., practice) across both halves of the task (first half: β = .088, *SE* = .013, *p* < .001; second half: β = .121, *SE* = .013, *p* < .001). There was no independent effect of wave on network modularity. This suggests that while modularity tends to increase within-session across the task and with older individuals, there is no independent effect of repeated exposure to the same task environment (Table 3). As expected, this model offered improvements over the random effects ANOVA model 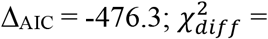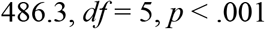.

#### 3.3.2 Network Modularity Shows Interactions Across Levels of Time

We next fit the interactive model of development to the network modularity data. Similar to the model with learning performance, there was a significant interaction of wave and the quadratic effect of age on network modularity (β =.057, *SE* = .018, *p* = .001; Table 4). When probed (Figure 6A), there was a similar compensatory pattern to the one seen in learning performance, such that experience across waves (green and blue) predicted positive shifts in modularity at later ages compared with the first exposure to the task (red). Interestingly, these differences appear to only emerge during the transition from adolescence to young adulthood, whereas increased experience does not impact modularity at younger ages. Furthermore, there was a significant positive interaction of age and practice in the first half of the task (β = .068, *SE* = .019, *p* < .001), such that older individuals showed more rapid gains in modularity across the first half of the task (Figure 6B). This model offered continued improvements over the main effects only model 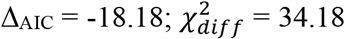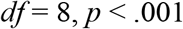.

**Figure 6.**
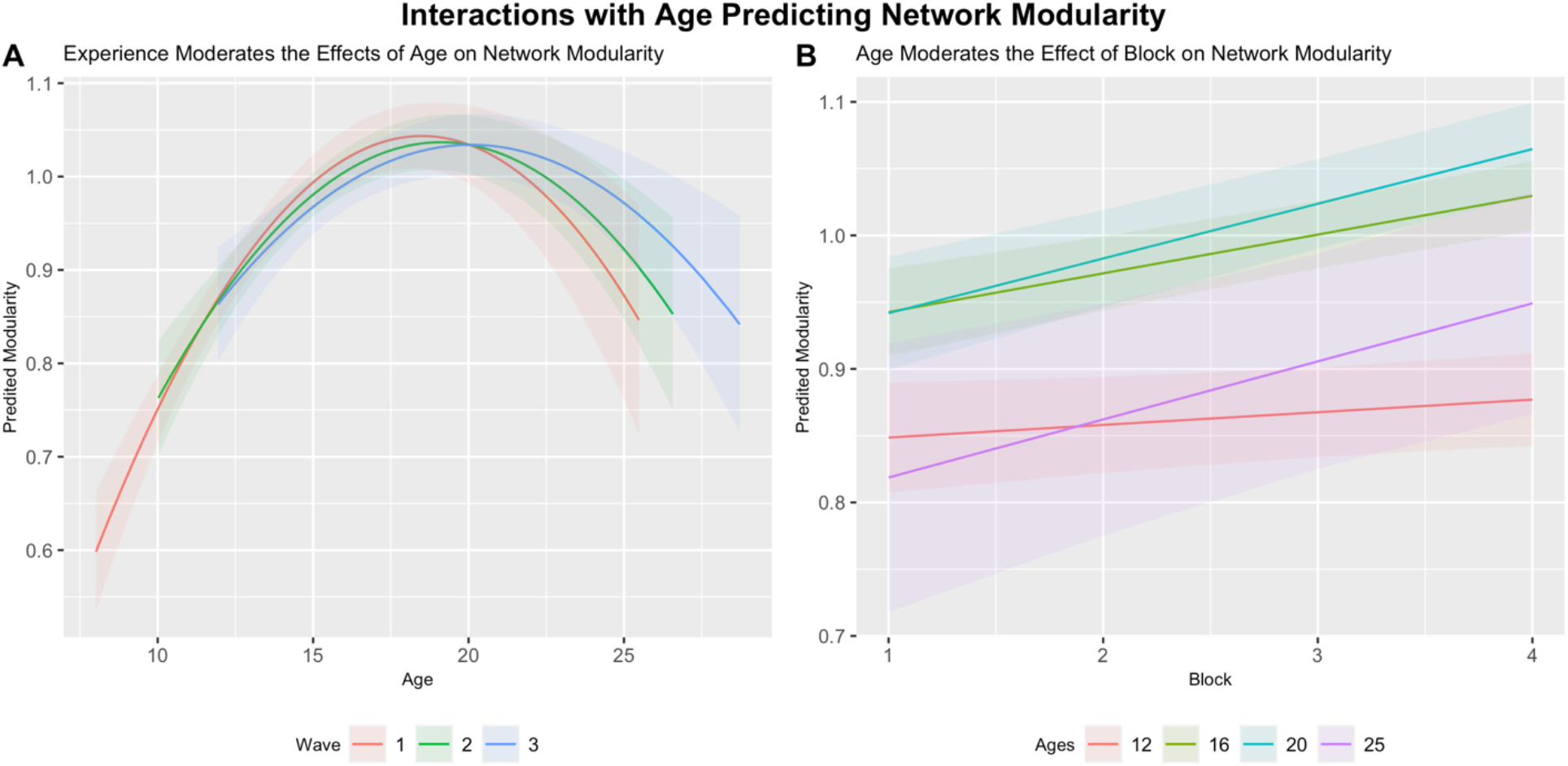
Experience and Age Impact Brain Network Organization. A) Accumulating experience (green and blue) across waves predicts increased network modularity compared with first exposure (red), however these differences only emerge during the transition from adolescence to young adulthood. B) Increased age predicts greater positive gains in network modularity across blocks in the first half of the task.

#### 3.3.3 Predicting Learning with Network Modularity

Finally, we tested whether network modularity predicts learning performance above and beyond the effects of time. To do so, we entered network modularity and interaction terms between modularity and the time predictors into the model. In addition to similar effects of the time predictors, this model revealed a significant positive interaction between network modularity and age (β = .062, *SE* = .023, *p* = .007; Table 5). Probing this interaction (Figure 7), we plotted the model-implied trajectory of learning rate across age for individuals who showed relatively low (−1 SD), mean, and relatively high (+1 SD) network modularity. This showed that individuals who evinced relatively high network modularity (blue trajectory) tended to show higher learning rates, but only after the transition from adolescence to young adulthood (see region of significance analysis in Supplement for details). A likelihood ratio test suggests that including the brain as a predictor increased model fit 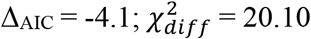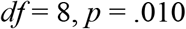.

**Figure 7.**
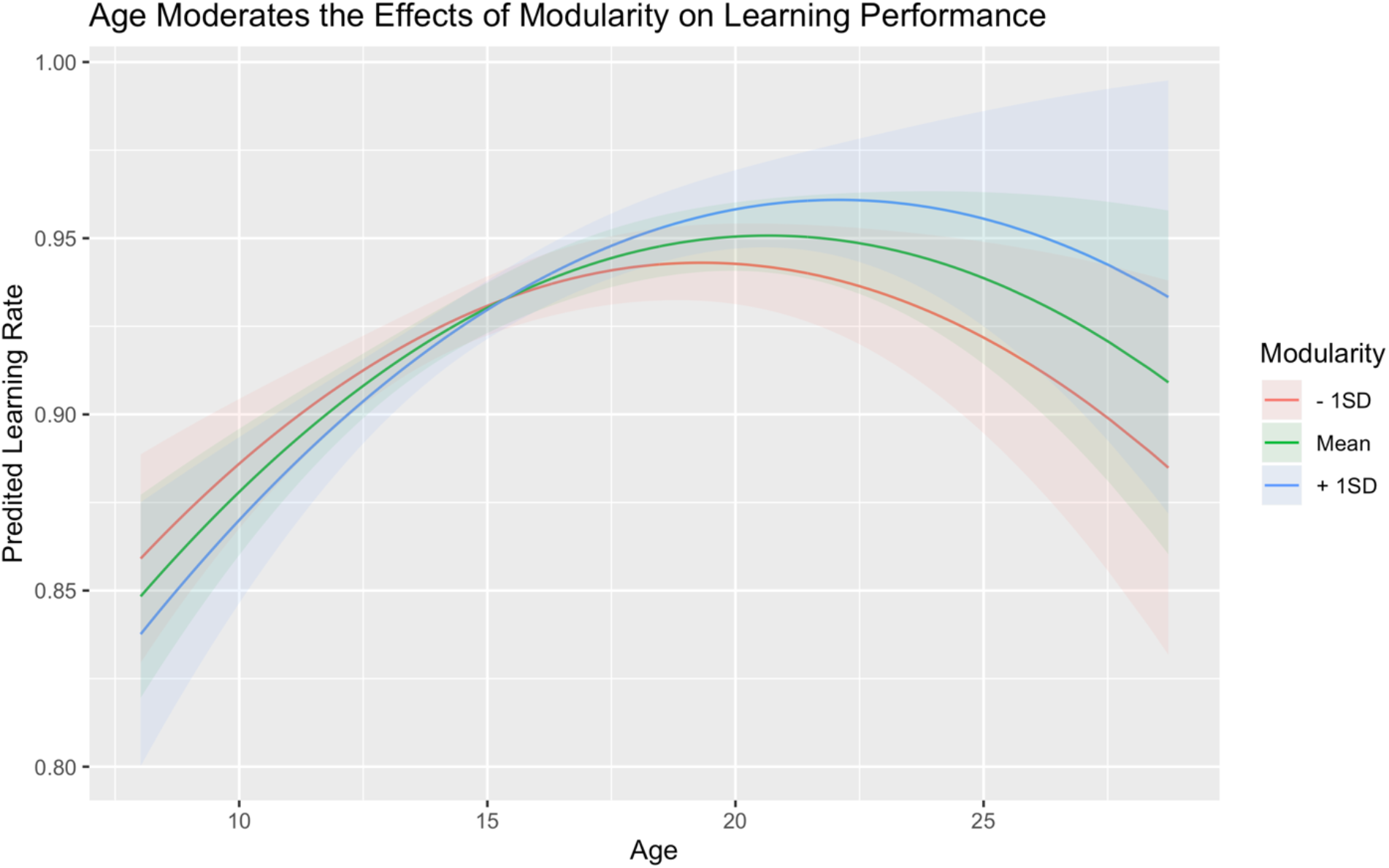
Differential Impacts of Network Modularity on Learning Rate By Age. At earlier ages, increased modularity did not predict increased learning, but this relationship emerges following adolescence.

**Table 5.**
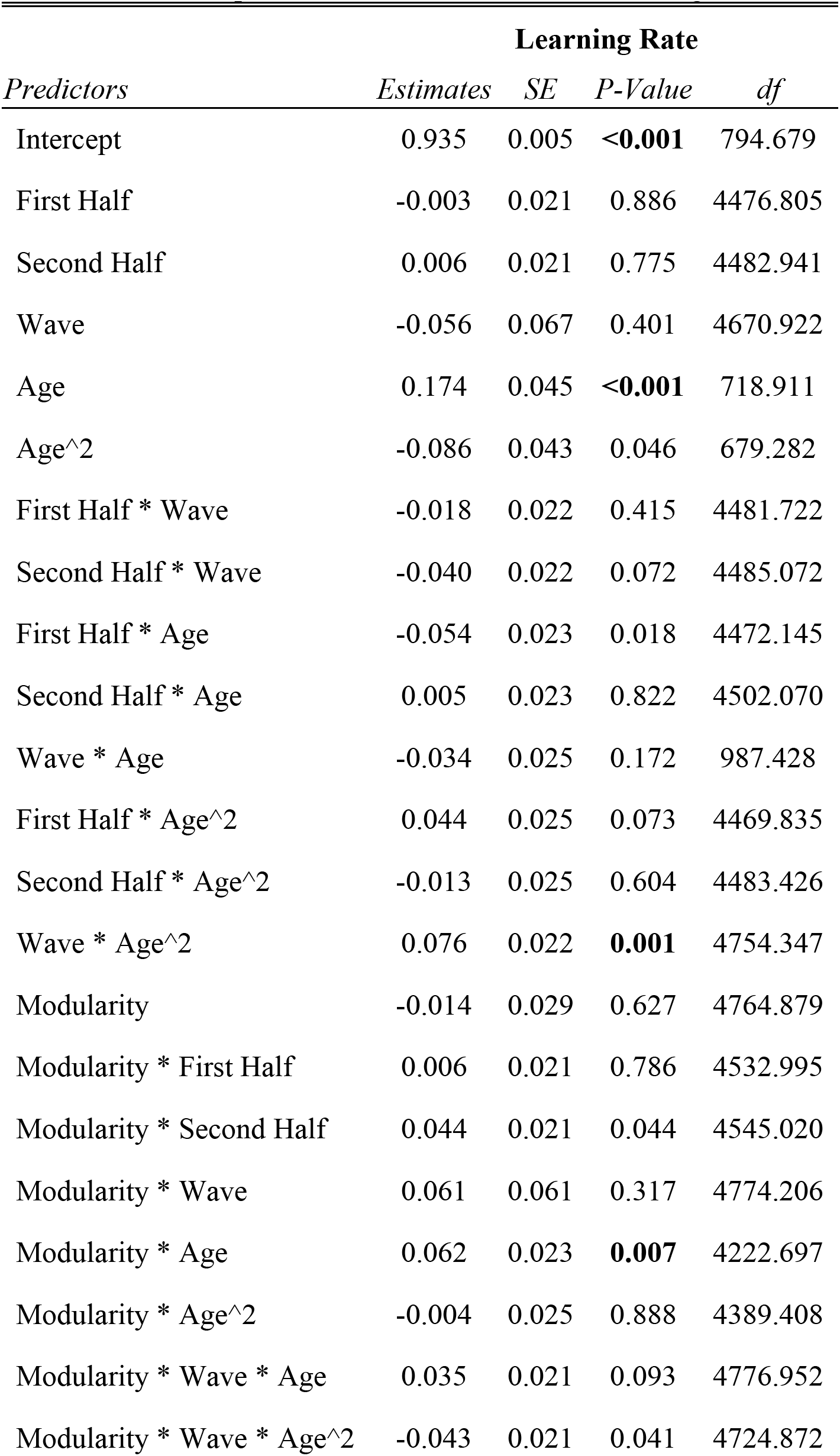
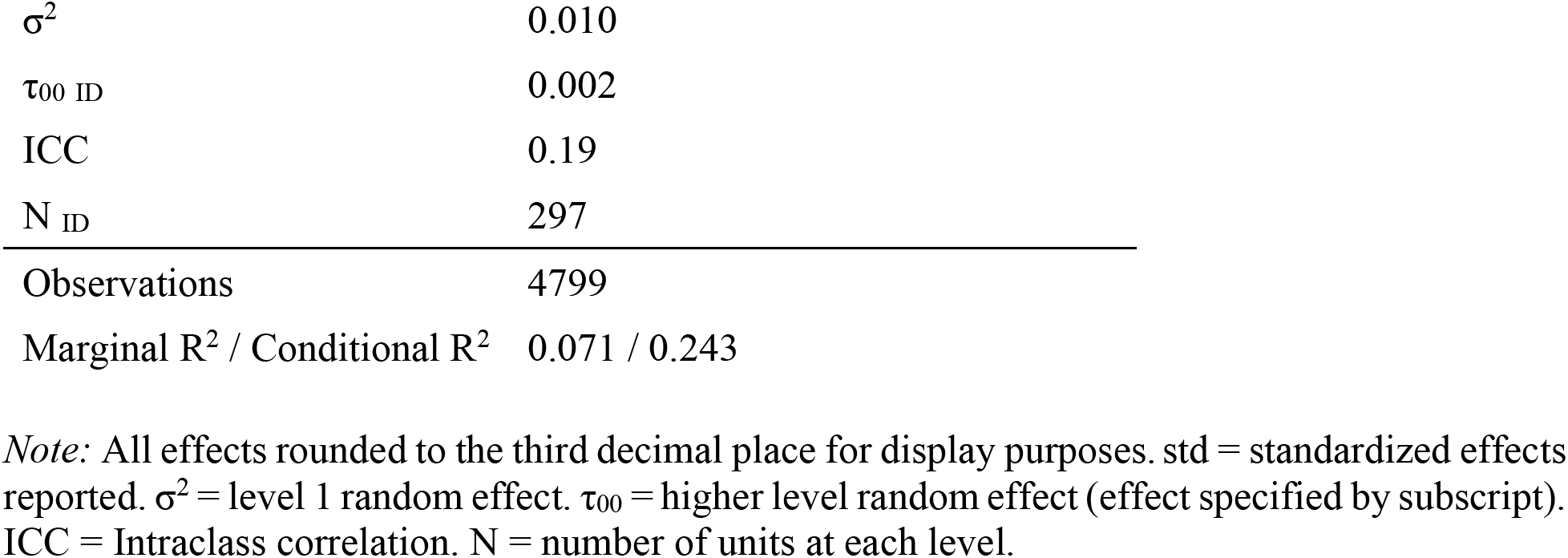
Model Output from Brain as Predictor of Learning Performance Model.

| *Predictors* | **Learning Rate** |  |  |  |
| --- | --- | --- | --- | --- |
|  | *Estimates* | *SE* | *P-Value* | *df* |
| Intercept | 0.935 | 0.005 | **<0.001** | 794.679 |
| First Half | -0.003 | 0.021 | 0.886 | 4476.805 |
| Second Half | 0.006 | 0.021 | 0.775 | 4482.941 |
| Wave | -0.056 | 0.067 | 0.401 | 4670.922 |
| Age | 0.174 | 0.045 | **<0.001** | 718.911 |
| Age^2 | -0.086 | 0.043 | 0.046 | 679.282 |
| First Half * Wave | -0.018 | 0.022 | 0.415 | 4481.722 |
| Second Half * Wave | -0.040 | 0.022 | 0.072 | 4485.072 |
| First Half * Age | -0.054 | 0.023 | 0.018 | 4472.145 |
| Second Half * Age | 0.005 | 0.023 | 0.822 | 4502.070 |
| Wave * Age | -0.034 | 0.025 | 0.172 | 987.428 |
| First Half * Age^2 | 0.044 | 0.025 | 0.073 | 4469.835 |
| Second Half * Age^2 | -0.013 | 0.025 | 0.604 | 4483.426 |
| Wave * Age^2 | 0.076 | 0.022 | **0.001** | 4754.347 |
| Modularity | -0.014 | 0.029 | 0.627 | 4764.879 |
| Modularity * First Half | 0.006 | 0.021 | 0.786 | 4532.995 |
| Modularity * Second Half | 0.044 | 0.021 | 0.044 | 4545.020 |
| Modularity * Wave | 0.061 | 0.061 | 0.317 | 4774.206 |
| Modularity * Age | 0.062 | 0.023 | **0.007** | 4222.697 |
| Modularity * Age^2 | -0.004 | 0.025 | 0.888 | 4389.408 |
| Modularity * Wave * Age | 0.035 | 0.021 | 0.093 | 4776.952 |
| Modularity * Wave * Age^2 | -0.043 | 0.021 | 0.041 | 4724.872 |

**Random Effects**
|  |  |
| --- | --- |
| $\sigma^2$ | 0.010 |
| $\tau_{00 \text{ ID}}$ | 0.002 |
| ICC | 0.19 |
| $N_{\text{ID}}$ | 297 |
| Observations | 4799 |
| Marginal $R^2$ / Conditional $R^2$ | 0.071 / 0.243 |

*Note:* All effects rounded to the third decimal place for display purposes. std = standardized effects reported. $\sigma^2$ = level 1 random effect. $\tau_{00}$ = higher level random effect (effect specified by subscript). ICC = Intraclass correlation. N = number of units at each level.

## 4. Discussion

The contributions of experience versus development can be difficult to tease apart, especially within a longitudinal sample (Jolles & Crone, 2012; Telzer et al., 2018) because age, experience, and practice are confounded. To test an interaction model of experience and development, we utilized a novel longitudinal approach to separate out variability in learning performance and brain network modularity along different timescales: 1) within-session practice across blocks of learning, 2) within-person, across-waves, and 3) across age. Briefly, we found that learning performance tends to improve throughout adolescence and level off into adulthood. In contrast, network modularity appears to peak around middle adolescence and then shows declines across young adulthood. However, both of these effects are moderated by the amount of experience individuals have accumulated across waves, with more experience relating to higher performance and network modularity during learning. When considering the brain-behavior relationships, relatively higher modularity predicts better learning performance but only at older ages, suggesting that the importance of individual differences around developmental trends for determining behavioral outcomes may depend on timing in development. We discuss each of these findings in greater detail below.

### 4.1 Multi-Level Changes in Learning Rate and Brain Networks

Even without probing cross-level interactions, a major advantage of the models employed here is the separation of variance in the outcome across levels of time. In the current study, we found that behavioral improvements in learning occur across age, but do not change systematically across waves or blocks within the task. In contrast, network modularity showed systematic within-session increases as well as positive age effects, while showing no systematic changes across waves. While these total effects should be interpreted with caution given the presence of higher-order interaction effects, they nevertheless highlight an advantage of the growth model employed here (McCormick, *preprint*). Controlling for the effects of repeated exposure to the task environment allow us to be more confident that the effects of age are due to maturational forces rather than comfort or familiarity with the task or scanner environment (Bell, 1953; Jolles & Crone, 2012). Even if these effects are not significant, as is the case here, these models allow us to check our assumptions about the processes underlying longitudinal change.

### 4.2 Cross-Level Interactions between Growth at Different Scales

More interestingly, there were significant cross-level interactions between time predictors in the models for both behavior and brain trajectories. For both outcomes of interest, results supported the idea of an interactive model of development, where the impacts of experience change across age. For learning rate, the model recapitulated previous findings in this task (Peters & Crone, 2017) showing that learning rate reaches peak levels around late adolescence, and is consistent with other findings showing improvements in learning across adolescence (van Duijvenvoorde et al., 2008; Peters et al., 2016; McCormick & Telzer, 2017a). This highlights that the model with growth at multiple levels accommodates the same inferences made with the age-only models. In this way, adopting the current approach offers advantages without limiting the inferences made about developmental trajectories. Indeed, the interaction of wave and the quadratic effect of age on learning performance reveals that changes in behavior are driven by complex interplays between development and experience, with experience appearing to play different roles in supporting overall performance across adolescence and young adulthood. Specifically, probing the interaction shows that increased experience compensates for the main effect of age, boosting behavioral performance gains further into to young adulthood. Extensive experience (blue trajectory in Figure 5) blunts otherwise predicted declines and stabilizes performance at the peak achieved during middle to late adolescence. However, this experience-related improvement is not universal across the developmental period considered, as mid-adolescent performance is heightened even at first wave. This might suggest that during adolescence, learning is universally improved, and then individual differences become more relevant as individuals transition out of this period (see Pattwell et al., 2011 for a similar idea in the area of contextual fear during adolescence). None of the models tested showed within-session effects on learning performance, although we may have been under-powered to detect such effects given the restraints of the task (i.e., there was a maximum of 12 trials per block regardless of learning). Overall, these results are a powerful validation of the model, converging with previous findings on the same data while simultaneously allowing for the addition of more complex time-dependent relationships.

Findings with the brain showed a distinct pattern of effects from growth in learning performance. Previous work in this area has shown developmental changes during learning in regional activation of the fronto-parietal (Peters et al., 2016) and striatal regions (Peters & Crone, 2017; McCormick & Telzer, 2017a), as well as seed-based connectivity with the orbitofrontal cortex (McCormick & Telzer, 2017a). In the current study, we instead examine changes in brain network organization during learning. Learning over short periods of time has been shown to alter neural network organization (Bassett et al., 2011; Telesford et al., 2017; Gerraty et al., 2018), but these processes have not been compared with long-term changes due to maturation previously. In the current study, we show changes in network modularity both at the short-term within-session and long-term developmental level. In the short term, playing repeated blocks within-session is associated with heightened modularity between brain networks involved in learning, and this effect across the first half of the task increases with age. This suggests that older individuals are able to segregate (i.e., high within-network and low between-network connectivity) the relevant networks of regions to a greater extent than younger participants. The quadratic total effect of age suggests that rather than a linear increase in modularity across experience across waves and maturation, network modularity instead decreases at later ages (Figure 6A), with late adolescents and young adults showing decreasingly modular networks compared with middle-adolescents (also consistent with the age-only model (Figure 4D). This pattern of network connectivity evolves similarly across development to striatal activation during learning versus application (Peters & Crone, 2017). However, the interaction of this age effect with wave shows an analogous effect as seen in learning performance, where experience appears to blunt expected decreases in network modularity in young adulthood. While this compensatory effect does not counteract the overall decreases as seen in the behavioral results, it does serve to shift network organization toward a more adolescent-typical phenotype. For both behavioral performance and network modularity, the pattern of results seen here clearly support an interactive view of experience and development, where maturation constrains the effect of experience rather than simply serving as a form of extended practice.

Regardless of the effects of experience (i.e., wave), there appear to be more substantial decreases in modularity compared with behavioral performance. In the context of learning and development, this pattern of long-term change in neural networks might have two potential explanations. First, it might be that brain networks show the greatest capacity for modularity during adolescence, and this capacity supports the heightened flexible learning (Johnson & Wilbrecht, 2011; Casey, 2015) and feedback sensitivity (Peters et al., 2016; van Duijvenvoorde et al., 2014; McCormick & Telzer, 2017a; b; 2018b) that characterize this developmental period regardless of practice or exposure effects. Alternatively, these changes might reflect differences in how the brain performs similar actions across development. For instance, high modularity might be necessary for adolescents to achieve high performance, while adults might not require this to the same degree. For instance, stabilizing learning performance with experience is still associated with overall (albeit blunted) decreases in modularity during young adulthood. This might also be consistent with an expansion-normalization theory of development and learning (Wenger et al., 2017) where brains show initial changes in structure and function (e.g., increased synaptic formation, increased activation) that then return to baseline without compromising behavioral performance. These hypotheses are not mutually exclusive, and the effects probed here might suggest both that adolescent brains support improved performance regardless of other influences (e.g., experience) and that returns to network phenotypes seen in younger adolescents does not lead to a complete collapse of behavioral performance.

### 4.3 Changing Brain-Behavior Relationships

While characterizing trajectories of brain and behavior separately is informative, we also examined whether individual differences in network organization might predict learning performance above and beyond developmental (Peters & Crone, 2017) and experiential effects, as well as whether that relationship changed across time. Consistent with the interaction view of development and experience, significant interactions between network modularity and time predictors in the model revealed a significant moderation of brain-behavior relationships across age. This means that in early adolescence, there is no effect of increased modularity on learning performance, but that increased modularity predicts enhanced learning rates for older individuals. These findings are consistent with previous work in young adults showing positive associations between network modularity and successful learning (Bassett et al., 2011; Ellefsen et al., 2015) and higher-order cognitive processing generally (Kitzbichler et al., 2011; Braun et al., 2015). However, these positive associations at later ages are particularly interesting in the context of the developmental trends detected, where older individuals on average show decreased modularity across waves. Despite normative decreases in network modularity during late adolescence and young adulthood, individuals with higher modularity show greater learning performance. This might lend support to the first explanation of the interaction effect of age and wave – that adolescent-typical neural phenotypes offer advantages in performance (e.g., Johnson & Wilbrecht, 2011; Jones et al., 2014; van Duijvenvoorde et al., 2016). If young adults simply did not need highly modular networks to perform what is a relatively easy task for them (i.e., the expansion-renormalization hypothesis), then we would expect that the relationship between modularity and learning performance would decrease. However, the positive interaction effect (Figure 7) suggests that older individuals show an even greater dependence on high modularity for successful learning performance compared with younger participants. This pattern of results presents a compelling case for adolescent-specific advantages in learning. For older individuals, retaining “immature,” but apparently more-optimal, network configurations helps boost behavioral performance.

### 4.4 Conclusions

In summary, we took a novel modeling approach to disaggregate change in learning and neural networks across different timescales. While we focus on learning here, these results highlight the potential flexibility of mixed-effects models for probing complex developmental trajectories in other domains. We found nonlinear patterns of development for both behavior and brain, which were moderated by experience. Specifically, greater experience with the task across waves supported increased learning and network modularity relative to comparatively naïve subjects at later ages, highlighting that these effects can potentially bias age-related inferences unless explicitly included in longitudinal models. Future research using accelerated longitudinal designs (or see McCormick, *preprint* for alternatives) should take care to model practice/exposure-related effects to remove this confound. Finally, we showed changing brain-behavior relationships across adolescence, where higher network modularity predicts increased learning performance only following the transition into young-adulthood. These results present compelling support for an interactive view of experience and development, where changes in the brain impact behavior in context-specific fashion based on developmental goals (Crone & Dahl, 2012; Romer et al., 2017).

## Acknowledgements

Author Contributions: S.P. and E.A.C. designed research, and performed research; E.M.M., analyzed data; E.M.M., S.P., E.A.C. & E.H.T. wrote the paper.

This research was funded by a starting grant of the European Research Council (ERC-2010-StG-263234 awarded to E.A.C.) and a grant from the Netherlands Organization for Scientific Research (NWO-VICI 453-14-001 awarded to E.A.C.). We would like to thank Laura van der Aar, Sibel Altikulaç, Neeltje Blankenstein, Barbara Braams, Suzanne van de Groep, Juliette Cassé, Dianne van der Heide, Jorien van Hoorn, Cédric Koolschijn, Babette Langeveld, Kyra Lubbers, Batsheva Mannheim, Mara van der Meulen, Rosa Meuwese, Sandy Overgaauw, Jiska Peper, Elisabeth Schreuders, Merel Schrijver, Jochem Spaans, Marije Stolte, Erik de Water, and Bianca Westhoff for their help with data collection, Ferdi van de Kamp for help with data analyses, and Anna Van Duijvenvoorde and Berna Güroğlu for carefully reviewing the manuscript. Finally we would like to thank all participants and their parents for their collaboration.

E.M.M. was supported in this research by grants from the National Institutes of Health (R01DA039923, R01 EB022904 awarded to E.H.T.) and generous funds from the University of North Carolina at Chapel Hill.

The authors declare no competing financial interests.

