## Supplemental Figure and Code for "Longitudinal Network Re-organization Across Learning and Development"

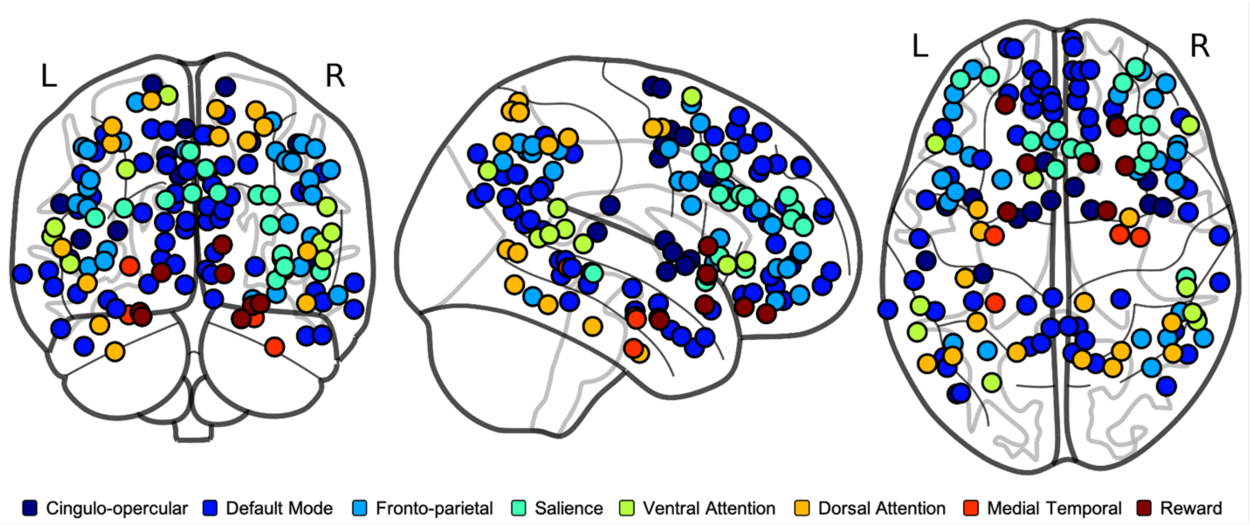


*Figure S1. Coordinates locations of ROIs used to compute network modularity. Distinct sub-networks are shown by unique colors.*

### BrainTime Project

Ethan McCormick

1/15/2020

#### Read in Data and Create Centered Variables

```
## Read in Data
BTdat = read.csv('~/Dropbox/Professional/Research_Projects/BrainTime/braintime_mlmdata.csv', header = T)
output.dir = '/Users/Ethan/Dropbox/Professional/Research_Projects/BrainTime/_plots/'

## Create Centered Variables
wave_c = BTdat$wave - 2
age_c = scale(BTdat$age, center=TRUE, scale=FALSE)
modrescale = scale(BTdat$modavg, center=FALSE, scale=TRUE)
modrescale_c = scale(modrescale, center=TRUE, scale=FALSE)
BTdat = cbind(BTdat, wave_c, age_c, modrescale, modrescale_c)

## Specify Interaction Levels to Probe
int.ages = c(12,16,20,25) - mean(BTdat$age)
int.ages
```

```
## [1] -3.8536424  0.1463576  4.1463576  9.1463576
```

#### Descriptives & Sample Distributions

Age

| wave | M | SD | min | max |
|---|---|---|---|---|
| 1 | 14.16771 | 3.631478 | 8.01 | 25.95 |
| 2 | 16.14764 | 3.618389 | 10.02 | 26.61 |
| 3 | 18.14819 | 3.682511 | 11.94 | 28.72 |

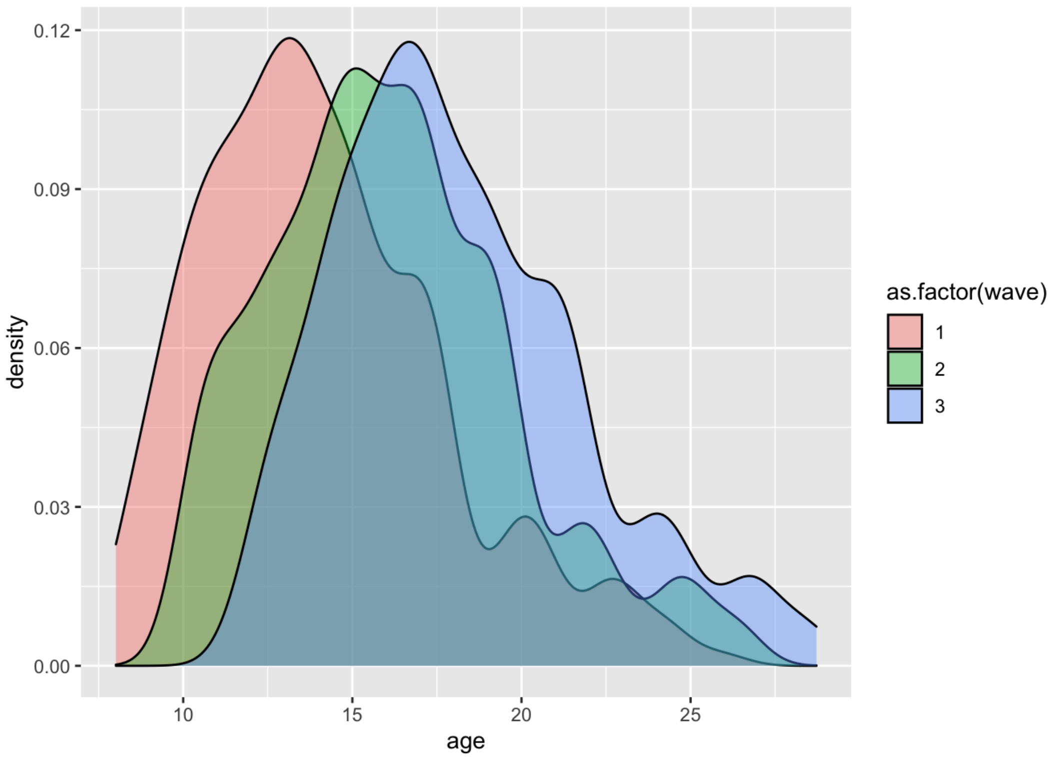

Pubertal Development

| wave | M | SD | min | max |
|---|---|---|---|---|
| 1 | 2.238998 | 0.9283809 | 0.4 | 4 |
| 2 | 2.596512 | 0.8644957 | 0.6 | 4 |
| 3 | 2.922835 | 0.7227913 | 1.0 | 4 |

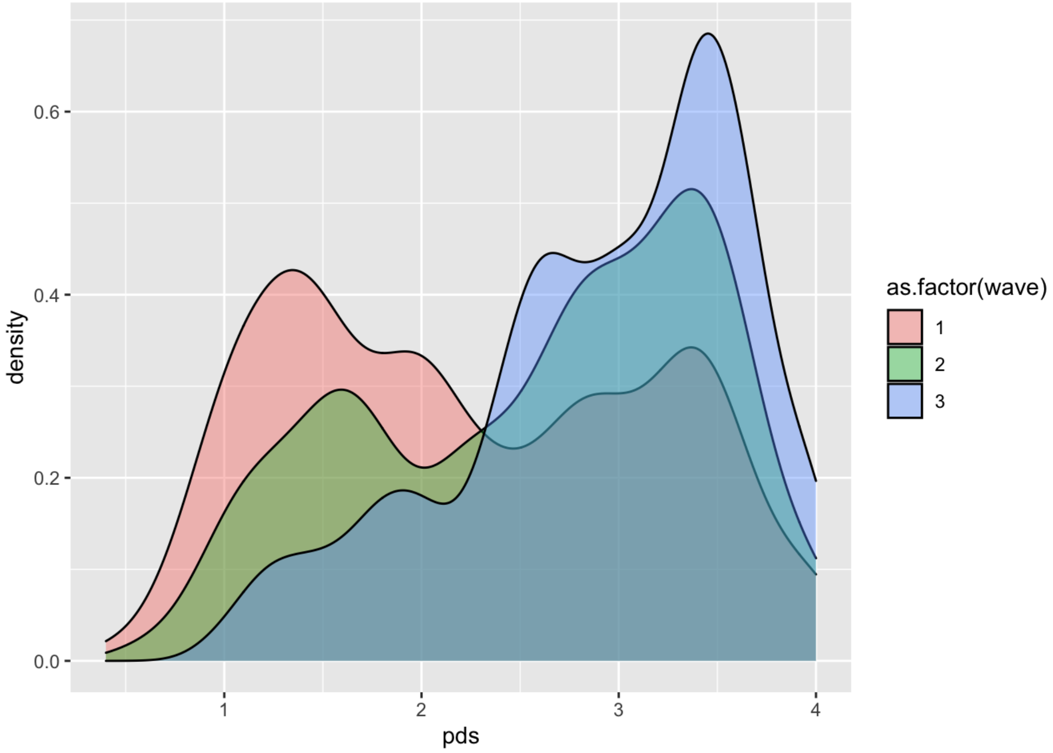


IQ

| wave | M | SD | min | max |
|---|---|---|---|---|
| 1 | 109.8185 | 10.29618 | 80 | 142.5 |
| 2 | 108.3402 | 10.26893 | 80 | 147.5 |
| 3 | NaN | NA | Inf | -Inf |

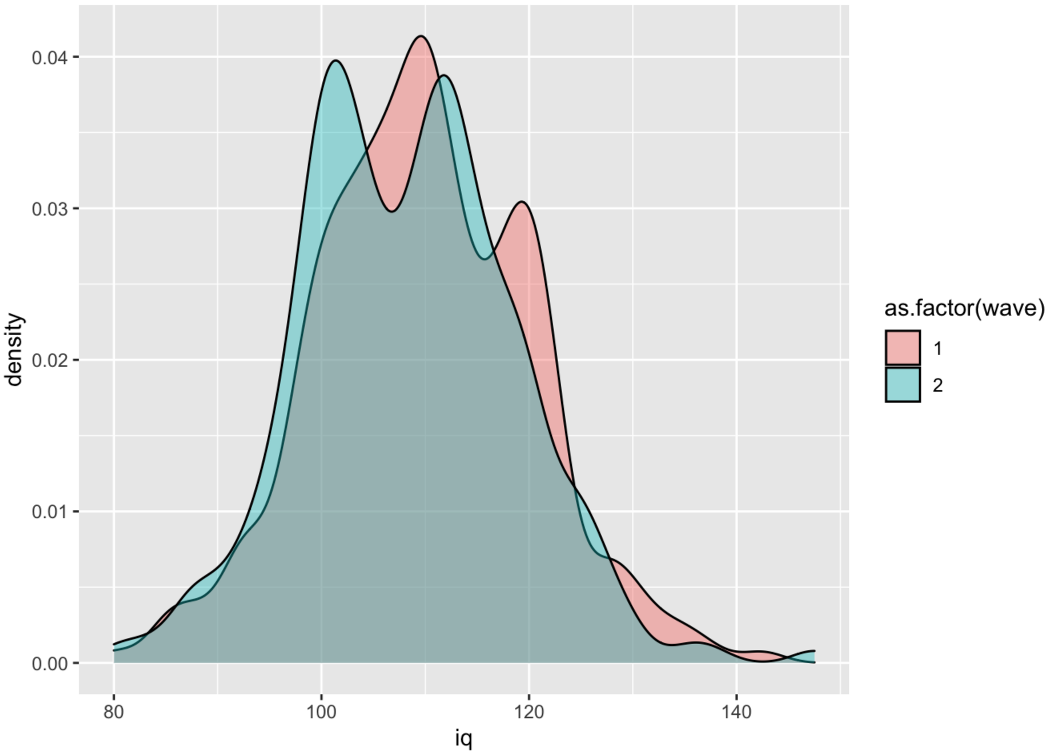


```
## # A tibble: 2 x 2
##    male     n
##   <int> <int>
## 1     0   153
## 2     1   144
```

```
## # A tibble: 6 x 3
## # Groups:   wave [3]
##    wave  male     n
##   <int> <int> <int>
## 1     1     0   140
## 2     1     1   134
## 3     2     0   121
## 4     2     1   121
## 5     3     0   122
## 6     3     1   116
```

### Random Effects ANOVA Models

Specify models with no predictors and only random effects at level 1 and level 2:

```
REAM.behav = lmer(learnrate  ~ 1 + (1 | id), data = BTdat)
REAM.brain = lmer(modrescale ~ 1 + (1 | id), data = BTdat)
```

| | Learning Rate | | | | Network Modularity | | | |
|---|---|---|---|---|---|---|---|---|
| *Predictors* | *Estimates* | *SE* | *P-Value* | *df* | *Estimates* | *SE* | *P-Value* | *df* |
| Intercept | 0.925 | 0.003 | **<0.001** | 271.214 | 0.946 | 0.012 | **<0.001** | 289.668 |
| **Random Effects** | | | | | | | | |
| $\sigma^2$ | 0.01 | | | | 0.06 | | | |
| $\tau_{00}$ | 0.00 $_{id}$ | | | | 0.04 $_{id}$ | | | |
| ICC | 0.22 | | | | 0.40 | | | |
| N | 297 $_{id}$ | | | | 297 $_{id}$ | | | |
| Observations | 4799 | | | | 4799 | | | |

| | | |
|---|---|---|
| Marginal $R^2$ / Conditional $R^2$ | 0.000 / 0.218 | 0.000 / 0.398 |

Specify models with no predictors and only random effects at levels 1-3:

```
REAM3.behav = lmer(learnrate  ~ 1 + (1 | id:wave_c) + (1 | id), data = BTdat)
REAM3.brain = lmer(modrescale ~ 1 + (1 | id:wave_c) + (1 | id), data = BTdat)
```

| | Learning Rate | | | | Network Modularity | | | |
|---|---|---|---|---|---|---|---|---|
| *Predictors* | *Estimates* | *SE* | *P-Value* | *df* | *Estimates* | *SE* | *P-Value* | *df* |
| Intercept | 0.927 | 0.003 | **<0.001** | 256.681 | 0.949 | 0.012 | **<0.001** | 284.150 |
| **Random Effects** | | | | | | | | |
| $\sigma^2$ | 0.01 | | | | 0.05 | | | |
| $\tau_{00}$ | 0.00 id:wave_c | | | | 0.01 id:wave_c | | | |
| | 0.00 id | | | | 0.03 id | | | |
| ICC | 0.31 | | | | 0.49 | | | |
| N | 297 id | | | | 297 id | | | |
| | 3 wave_c | | | | 3 wave_c | | | |
| Observations | 4799 | | | | 4799 | | | |
| Marginal $R^2$ / Conditional $R^2$ | 0.000 / 0.313 | | | | 0.000 / 0.486 | | | |

The variance inflation for the behavioral and brain models is

```
##    age_c
## 1.186597
```

### Descriptives

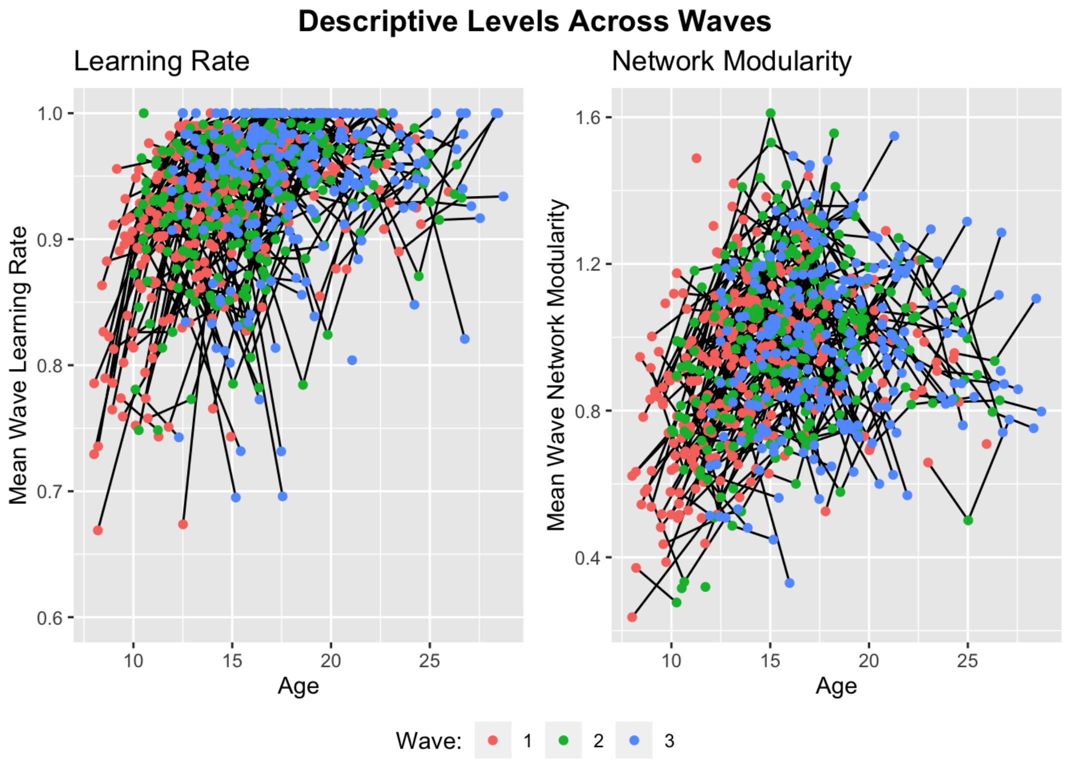


### Single Growth Models with Polynomials of Age

```
age.behav = lmer(learnrate ~ age_c + I(age_c^2) + (1 | id), data=BTdat)
age.brain = lmer(modrescale ~ age_c + I(age_c^2) + (1 | id), data=BTdat)
```

| | Learning Rate | | | | Network Modularity | | | |
|---|---|---|---|---|---|---|---|---|
| *Predictors* | *Estimates* | *SE* | *P-Value* | *df* | *Estimates* | *SE* | *P-Value* | *df* |
| Intercept | 0.937 | 0.004 | **<0.001** | 359.604 | 0.994 | 0.011 | **<0.001** | 361.688 |
| Age | 0.008 | 0.001 | **<0.001** | 1003.133 | 0.021 | 0.002 | **<0.001** | 1992.711 |
| Age^2 | -0.001 | 0.000 | **<0.001** | 1924.892 | -0.003 | 0.000 | **<0.001** | 3523.698 |
| **Random Effects** | | | | | | | | |
| $\sigma^2$ | 0.01 | | | | 0.05 | | | |
| $\tau_{00}$ | 0.00 $_{id}$ | | | | 0.03 $_{id}$ | | | |
| ICC | 0.18 | | | | 0.35 | | | |
| N | 297 $_{id}$ | | | | 297 $_{id}$ | | | |
| Observations | 4799 | | | | 4799 | | | |
| Marginal $R^2$ / Conditional $R^2$ | 0.065 / 0.236 | | | | 0.075 / 0.402 | | | |

```
##        term std.estimate  std.error   conf.low   conf.high
## 1      age_c    0.2774890 0.02287490  0.2326550  0.32232296
## 2 I(age_c^2)   -0.1396848 0.02056586 -0.1799931 -0.09937645
```

```
##        term std.estimate  std.error   conf.low  conf.high
## 1      age_c    0.2748599 0.02341206  0.2289732  0.3207467
## 2 I(age_c^2)   -0.2091728 0.01986656 -0.2481105 -0.1702351
```

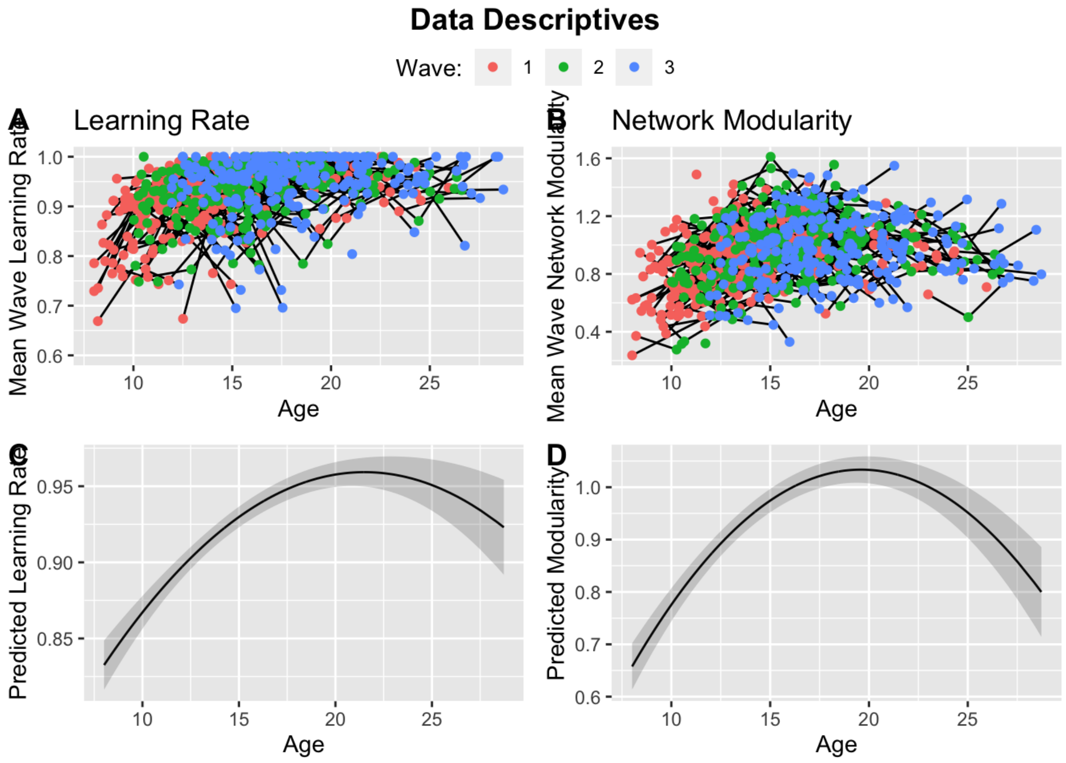



```
##        term std.estimate  std.error   conf.low   conf.high
## 1      age_c    0.2774890 0.02287490  0.2326550  0.32232296
## 2 I(age_c^2)   -0.1396848 0.02056586 -0.1799931 -0.09937645
```

```
##        term std.estimate  std.error   conf.low  conf.high
## 1      age_c    0.2748599 0.02341206  0.2289732  0.3207467
## 2 I(age_c^2)   -0.2091728 0.01986656 -0.2481105 -0.1702351
```

```
## JOHNSON-NEYMAN INTERVAL
##
## When age_c_temp is OUTSIDE the interval [8.61, 15.78], the slope of age_c
## is p < .05.
##
## Note: The range of observed values of age_c_temp is [-7.84, 12.87]
```

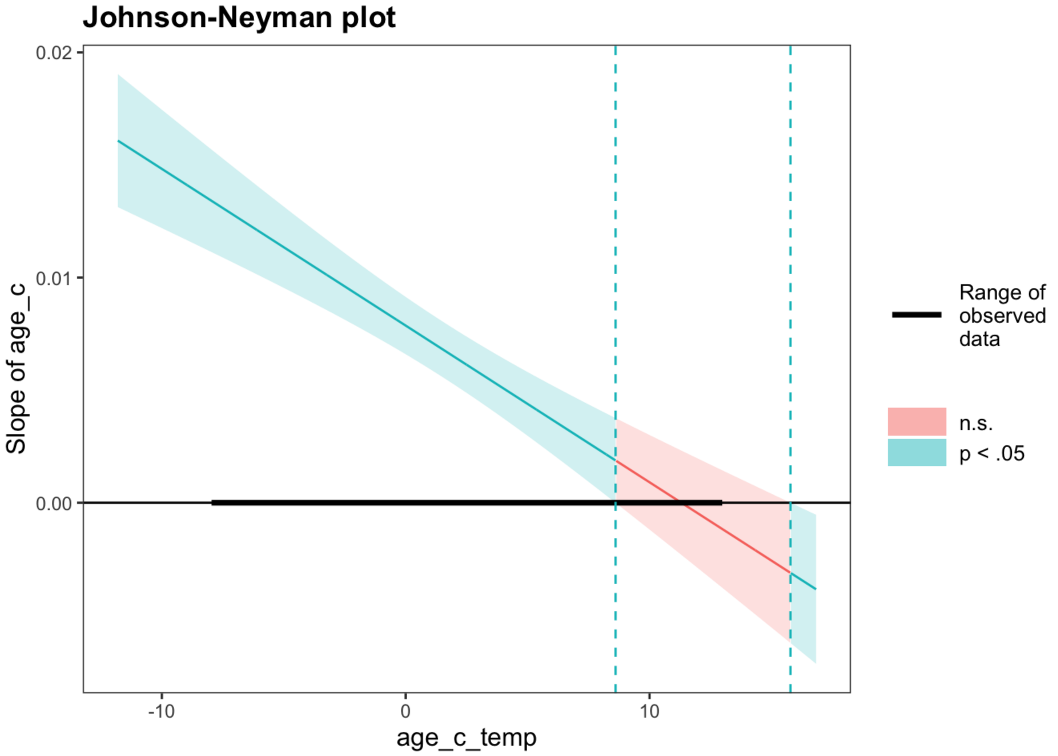


```
## SIMPLE SLOPES ANALYSIS
##
## Slope of age_c when age_c_temp = -3.96 (- 1 SD):
##
##   Est.   S.E.   t val.      p
## ------ ------ -------- ------
##   0.01   0.00    12.57   0.00
##
## Slope of age_c when age_c_temp =  0.00 (Mean):
##
##   Est.   S.E.   t val.      p
## ------ ------ -------- ------
##   0.01   0.00    12.13   0.00
##
## Slope of age_c when age_c_temp =  3.96 (+ 1 SD):
##
##   Est.   S.E.   t val.      p
## ------ ------ -------- ------
##   0.01   0.00     7.56   0.00
```

```
## JOHNSON-NEYMAN INTERVAL
##
## When age_c_temp is OUTSIDE the interval [5.97, 9.44], the slope of age_c is
## p < .05.
##
## Note: The range of observed values of age_c_temp is [-7.84, 12.87]
```

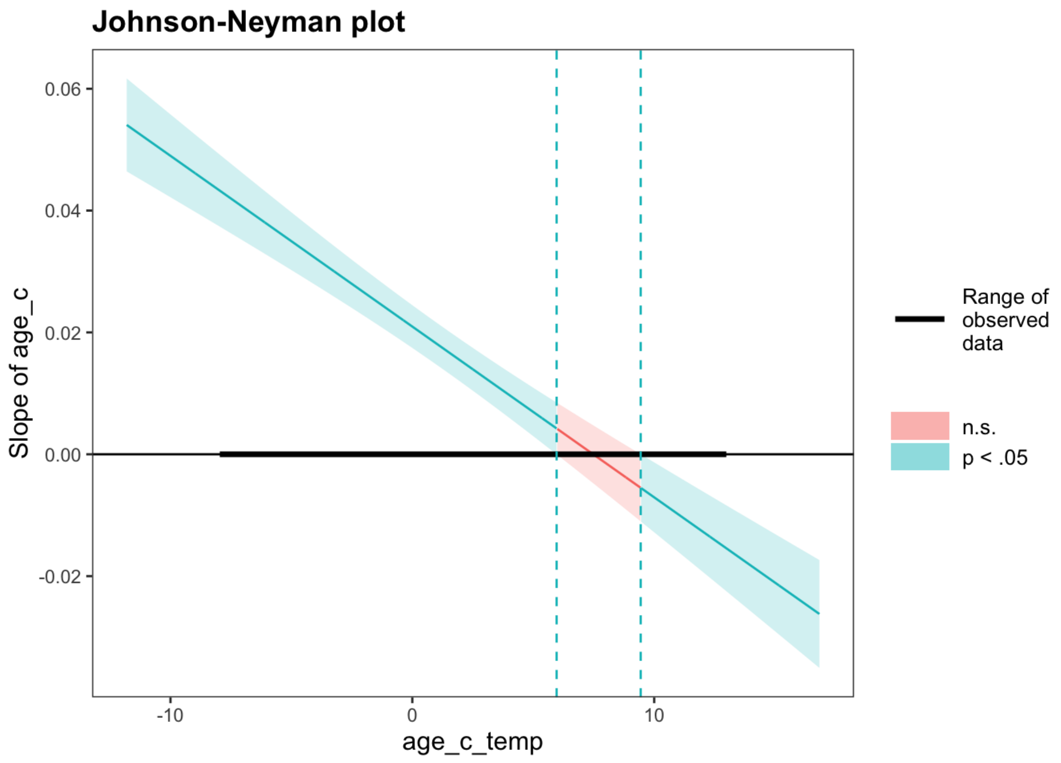


```
## SIMPLE SLOPES ANALYSIS
##
## Slope of age_c when age_c_temp = -3.96 (- 1 SD):
##
##   Est.   S.E.   t val.      p
## ------ ------ -------- ------
##   0.03   0.00    14.36   0.00
##
## Slope of age_c when age_c_temp =  0.00 (Mean):
##
##   Est.   S.E.   t val.      p
## ------ ------ -------- ------
##   0.02   0.00    11.74   0.00
##
## Slope of age_c when age_c_temp =  3.96 (+ 1 SD):
##
##   Est.   S.E.   t val.      p
## ------ ------ -------- ------
##   0.01   0.00     5.18   0.00
```

### Main Effects-Only Models

Specify models with linear independent predictors.

```
main.effects.behav = lmer(learnrate  ~ fh + sh + wave_c + age_c + I(age_c^2) + (1 | id), data = BTdat)
main.effects.brain = lmer(modrescale ~ fh + sh + wave_c + age_c + I(age_c^2) + (1 | id), data = BTdat)
```

| | Learning Rate | | | | Network Modularity | | | |
|---|---|---|---|---|---|---|---|---|
| *Predictors* | *Estimates* | *SE* | *P-Value* | *df* | *Estimates* | *SE* | *P-Value* | *df* |
| Intercept | 0.937 | 0.004 | **<0.001** | 717.508 | 0.987 | 0.013 | **<0.001** | 520.937 |
| First Half | 0.003 | 0.002 | 0.113 | 4473.435 | 0.025 | 0.004 | **<0.001** | 4491.297 |
| Second Half | 0.002 | 0.002 | 0.151 | 4473.468 | 0.035 | 0.004 | **<0.001** | 4491.319 |
| Wave | 0.003 | 0.003 | 0.198 | 1205.547 | -0.001 | 0.007 | 0.845 | 694.861 |
| Age | 0.007 | 0.001 | **<0.001** | 299.628 | 0.022 | 0.003 | **<0.001** | 311.080 |
| Age^2 | -0.001 | 0.000 | **<0.001** | 1941.031 | -0.003 | 0.000 | **<0.001** | 3747.429 |

| Random Effects | | |
|---|---|---|
| $\sigma^2$ | 0.01 | 0.05 |
| $\tau_{00}$ | 0.00 id | 0.03 id |
| ICC | 0.18 | 0.37 |
| N | 297 id | 297 id |
| Observations | 4799 | 4799 |
| Marginal $R^2$ / Conditional $R^2$ | 0.060 / 0.232 | 0.112 / 0.441 |

With standardized effects:

```
##         term std.estimate  std.error     conf.low   conf.high
## 1         fh   0.02479511 0.01565167 -0.005881595  0.05547181
## 2         sh   0.02247046 0.01564995 -0.008202877  0.05314380
## 3     wave_c   0.02329929 0.01809324 -0.012162800  0.05876139
## 4      age_c   0.24964394 0.03117515  0.188541776  0.31074611
## 5 I(age_c^2)  -0.13551763 0.02079267 -0.176270508 -0.09476476
```

```
##         term std.estimate  std.error    conf.low   conf.high
## 1         fh  0.088110844 0.01327006  0.06210200  0.11411969
## 2         sh  0.120916085 0.01326862  0.09491006  0.14692211
## 3     wave_c -0.003752798 0.01915563 -0.04129713  0.03379154
## 4      age_c  0.285223590 0.03894173  0.20889921  0.36154797
## 5 I(age_c^2) -0.208657572 0.01957416 -0.24702221 -0.17029293
```

Which offer a significant improvement over the random effects ANOVA model:

```
## refitting model(s) with ML (instead of REML)
```

```
## Data: BTdat
## Models:
## REAM.behav: learnrate ~ 1 + (1 | id)
## main.effects.behav: learnrate ~ fh + sh + wave_c + age_c + I(age_c^2) + (1 | id)
##                    Df     AIC     BIC logLik deviance  Chisq Chi Df Pr(>Chisq)
## REAM.behav          3 -7920.0 -7900.5 3963.0  -7926.0
## main.effects.behav  8 -8078.7 -8026.9 4047.3  -8094.7 168.71      5  < 2.2e-16
##
## REAM.behav
## main.effects.behav ***
## ---
## Signif. codes:  0 '***' 0.001 '**' 0.01 '*' 0.05 '.' 0.1 ' ' 1
```

```
## refitting model(s) with ML (instead of REML)
```

```
## Data: BTdat
## Models:
## REAM.brain: modrescale ~ 1 + (1 | id)
## main.effects.brain: modrescale ~ fh + sh + wave_c + age_c + I(age_c^2) + (1 | id)
##                    Df    AIC    BIC   logLik deviance Chisq Chi Df Pr(>Chisq)
## REAM.brain          3 550.79 570.22 -272.397   544.79
## main.effects.brain  8  74.49 126.30  -29.245    58.49 486.3      5  < 2.2e-16
##
## REAM.brain
## main.effects.brain ***
## ---
## Signif. codes:  0 '***' 0.001 '**' 0.01 '*' 0.05 '.' 0.1 ' ' 1
```

### Age-Levels Probed for Interactions with Respect to the Age Distribution

```
## Warning: Removed 2 rows containing missing values (geom_bar).
```

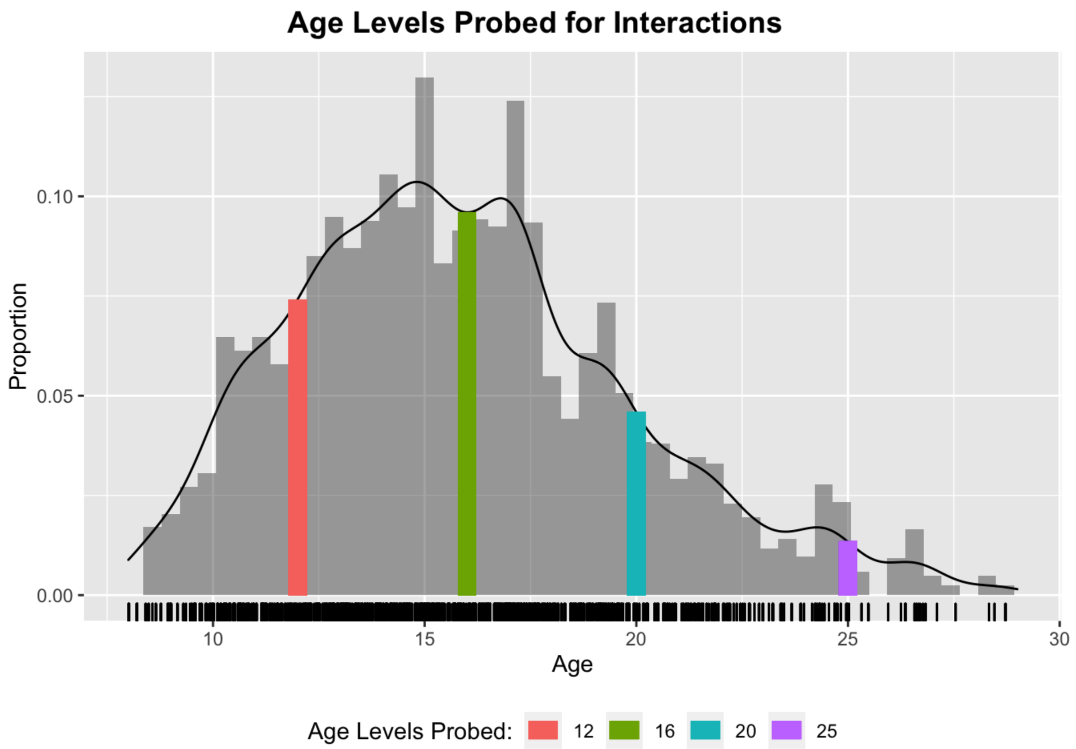


### Full Models with Interactions

```
int.behav = lmer(learnrate  ~ fh + sh + wave_c + age_c + I(age_c^2) +
                   fh:wave_c + sh:wave_c + fh:age_c + sh:age_c + wave_c:age_c +
                   fh:I(age_c^2) + sh:I(age_c^2) + wave_c:I(age_c^2) + (1 | id),
                 data = BTdat)

int.brain = lmer(modrescale  ~ fh + sh + wave_c + age_c + I(age_c^2) +
                   fh:wave_c + sh:wave_c + fh:age_c + sh:age_c + wave_c:age_c +
                   fh:I(age_c^2) + sh:I(age_c^2) + wave_c:I(age_c^2) + (1 | id),
                 data = BTdat)
```

| | **Learning Rate** | | | | **Network Modularity** | | | |
|---|---|---|---|---|---|---|---|---|
| *Predictors* | *Estimates* | *SE* | *P-Value* | *df* | *Estimates* | *SE* | *P-Value* | *df* |
| Intercept | 0.935 | 0.005 | **<0.001** | 768.901 | 0.997 | 0.014 | **<0.001** | 513.739 |
| First Half | -0.001 | 0.002 | 0.751 | 4473.838 | 0.027 | 0.005 | **<0.001** | 4495.715 |
| Second Half | 0.002 | 0.002 | 0.297 | 4473.787 | 0.033 | 0.005 | **<0.001** | 4495.776 |
| Wave | 0.002 | 0.004 | 0.722 | 3143.488 | -0.017 | 0.011 | 0.131 | 1734.974 |
| Age | 0.005 | 0.001 | **<0.001** | 650.191 | 0.025 | 0.004 | **<0.001** | 471.938 |
| Age^2 | -0.000 | 0.000 | 0.053 | 615.163 | -0.003 | 0.001 | **<0.001** | 513.285 |
| First Half * Wave | -0.002 | 0.002 | 0.469 | 4484.108 | -0.011 | 0.006 | 0.058 | 4508.580 |
| Second Half * Wave | -0.005 | 0.002 | **0.048** | 4484.488 | -0.007 | 0.006 | 0.244 | 4508.591 |
| First Half * Age | -0.001 | 0.000 | **0.027** | 4466.136 | 0.004 | 0.001 | **<0.001** | 4482.736 |
| Second Half * Age | 0.001 | 0.000 | 0.252 | 4466.143 | -0.000 | 0.001 | 0.818 | 4482.736 |
| Wave * Age | -0.001 | 0.001 | 0.111 | 870.145 | 0.000 | 0.003 | 0.947 | 648.001 |
| First Half * Age^2 | 0.000 | 0.000 | **0.048** | 4464.637 | -0.000 | 0.000 | 0.182 | 4481.364 |
| Second Half * Age^2 | -0.000 | 0.000 | 0.292 | 4464.630 | -0.000 | 0.000 | 0.906 | 4481.360 |

| | | | | | | | | |
|---|---|---|---|---|---|---|---|---|
| Wave * Age^2 | 0.000 | 0.000 | **<0.001** | 4746.733 | 0.001 | 0.000 | **0.001** | 4674.686 |
| **Random Effects** | | | | | | | | |
| $\sigma^2$ | 0.01 | | | | 0.05 | | | |
| $\tau_{00}$ | 0.00 $_{id}$ | | | | 0.03 $_{id}$ | | | |
| ICC | 0.19 | | | | 0.37 | | | |
| N | 297 $_{id}$ | | | | 297 $_{id}$ | | | |
| Observations | 4799 | | | | 4799 | | | |
| Marginal $R^2$ / Conditional $R^2$ | 0.066 / 0.239 | | | | 0.126 / 0.449 | | | |

Testing simple slopes:

```
## [redacted] While wave_c (2nd moderator) =  1.00 [redacted]
##
## JOHNSON-NEYMAN INTERVAL
##
## When age_c_temp is INSIDE the interval [-5.77, 9.31], the slope of age_c is
## p < .05.
##
## Note: The range of observed values of age_c_temp is [-7.84, 12.87]
##
## Interval calculated using false discovery rate adjusted t = 2.09
##
## SIMPLE SLOPES ANALYSIS
##
## Slope of age_c when age_c_temp = -3.96 (- 1 SD):
##
##   Est.   S.E.   t val.      p
## ------ ------ -------- ------
##   0.01   0.00     2.24   0.03
##
## Slope of age_c when age_c_temp =  0.00 (Mean):
##
##   Est.   S.E.   t val.      p
## ------ ------ -------- ------
##   0.00   0.00     2.68   0.01
##
## Slope of age_c when age_c_temp =  3.96 (+ 1 SD):
##
##   Est.   S.E.   t val.      p
## ------ ------ -------- ------
##   0.00   0.00     3.17   0.00
##
## [redacted] While wave_c (2nd moderator) =  0.00 [redacted]
##
## JOHNSON-NEYMAN INTERVAL
##
## When age_c_temp is OUTSIDE the interval [6.07, 21.47], the slope of age_c
## is p < .05.
##
## Note: The range of observed values of age_c_temp is [-7.84, 12.87]
##
## Interval calculated using false discovery rate adjusted t = 2.13
##
## SIMPLE SLOPES ANALYSIS
##
## Slope of age_c when age_c_temp = -3.96 (- 1 SD):
##
##   Est.   S.E.   t val.      p
## ------ ------ -------- ------
##   0.01   0.00     5.44   0.00
##
## Slope of age_c when age_c_temp =  0.00 (Mean):
##
##   Est.   S.E.   t val.      p
## ------ ------ -------- ------
```

```
##   0.01   0.00     5.64   0.00
##
## Slope of age_c when age_c_temp =  3.96 (+ 1 SD):
##
##   Est.   S.E.   t val.      p
## ------ ------ -------- ------
##   0.00   0.00     3.84   0.00
##
## [redacted] While wave_c (2nd moderator) = -1.00 [redacted]
##
## JOHNSON-NEYMAN INTERVAL
##
## When age_c_temp is OUTSIDE the interval [4.31, 13.65], the slope of age_c
## is p < .05.
##
## Note: The range of observed values of age_c_temp is [-7.84, 12.87]
##
## Interval calculated using false discovery rate adjusted t = 2.18
##
## SIMPLE SLOPES ANALYSIS
##
## Slope of age_c when age_c_temp = -3.96 (- 1 SD):
##
##   Est.   S.E.   t val.      p
## ------ ------ -------- ------
##   0.01   0.00     9.59   0.00
##
## Slope of age_c when age_c_temp =  0.00 (Mean):
##
##   Est.   S.E.   t val.      p
## ------ ------ -------- ------
##   0.01   0.00     7.52   0.00
##
## Slope of age_c when age_c_temp =  3.96 (+ 1 SD):
##
##   Est.   S.E.   t val.      p
## ------ ------ -------- ------
##   0.00   0.00     2.52   0.01
```

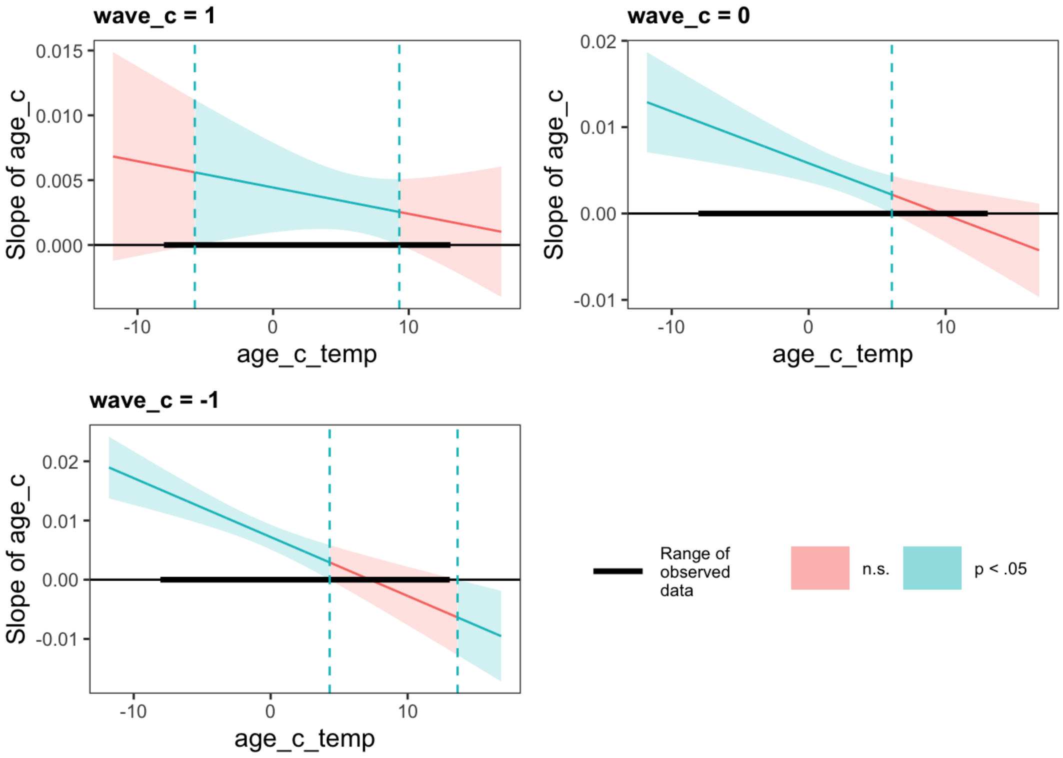


```
## [redacted] While wave_c (2nd moderator) =  1.00 [redacted]
##
## JOHNSON-NEYMAN INTERVAL
##
```

```
## When age_c_temp is OUTSIDE the interval [5.79, 12.48], the slope of age_c
## is p < .05.
##
## Note: The range of observed values of age_c_temp is [-7.84, 12.87]
##
## Interval calculated using false discovery rate adjusted t = 2.12
##
## SIMPLE SLOPES ANALYSIS
##
## Slope of age_c when age_c_temp = -3.96 (- 1 SD):
##
##   Est.   S.E.   t val.      p
## ------ ------ -------- ------
##   0.03   0.01     4.42   0.00
##
## Slope of age_c when age_c_temp =  0.00 (Mean):
##
##   Est.   S.E.   t val.      p
## ------ ------ -------- ------
##   0.02   0.01     4.23   0.00
##
## Slope of age_c when age_c_temp =  3.96 (+ 1 SD):
##
##   Est.   S.E.   t val.      p
## ------ ------ -------- ------
##   0.01   0.00     3.24   0.00
##
## [redacted] While wave_c (2nd moderator) =  0.00 [redacted]
##
## JOHNSON-NEYMAN INTERVAL
##
## When age_c_temp is OUTSIDE the interval [4.60, 9.49], the slope of age_c is
## p < .05.
##
## Note: The range of observed values of age_c_temp is [-7.84, 12.87]
##
## Interval calculated using false discovery rate adjusted t = 2.07
##
## SIMPLE SLOPES ANALYSIS
##
## Slope of age_c when age_c_temp = -3.96 (- 1 SD):
##
##   Est.   S.E.   t val.      p
## ------ ------ -------- ------
##   0.03   0.00     7.02   0.00
##
## Slope of age_c when age_c_temp =  0.00 (Mean):
##
##   Est.   S.E.   t val.      p
## ------ ------ -------- ------
##   0.02   0.00     6.34   0.00
##
## Slope of age_c when age_c_temp =  3.96 (+ 1 SD):
##
##   Est.   S.E.   t val.      p
## ------ ------ -------- ------
##   0.01   0.00     2.84   0.00
##
## [redacted] While wave_c (2nd moderator) = -1.00 [redacted]
##
## JOHNSON-NEYMAN INTERVAL
##
## When age_c_temp is OUTSIDE the interval [3.29, 8.63], the slope of age_c is
## p < .05.
##
## Note: The range of observed values of age_c_temp is [-7.84, 12.87]
##
## Interval calculated using false discovery rate adjusted t = 2.09
##
## SIMPLE SLOPES ANALYSIS
##
```

```
## Slope of age_c when age_c_temp = -3.96 (- 1 SD):
##
##   Est.   S.E.   t val.      p
## ------ ------ -------- ------
##   0.04   0.00    10.24   0.00
##
## Slope of age_c when age_c_temp =  0.00 (Mean):
##
##   Est.   S.E.   t val.      p
## ------ ------ -------- ------
##   0.02   0.00     6.96   0.00
##
## Slope of age_c when age_c_temp =  3.96 (+ 1 SD):
##
##   Est.   S.E.   t val.      p
## ------ ------ -------- ------
##   0.01   0.00     1.30   0.19
```

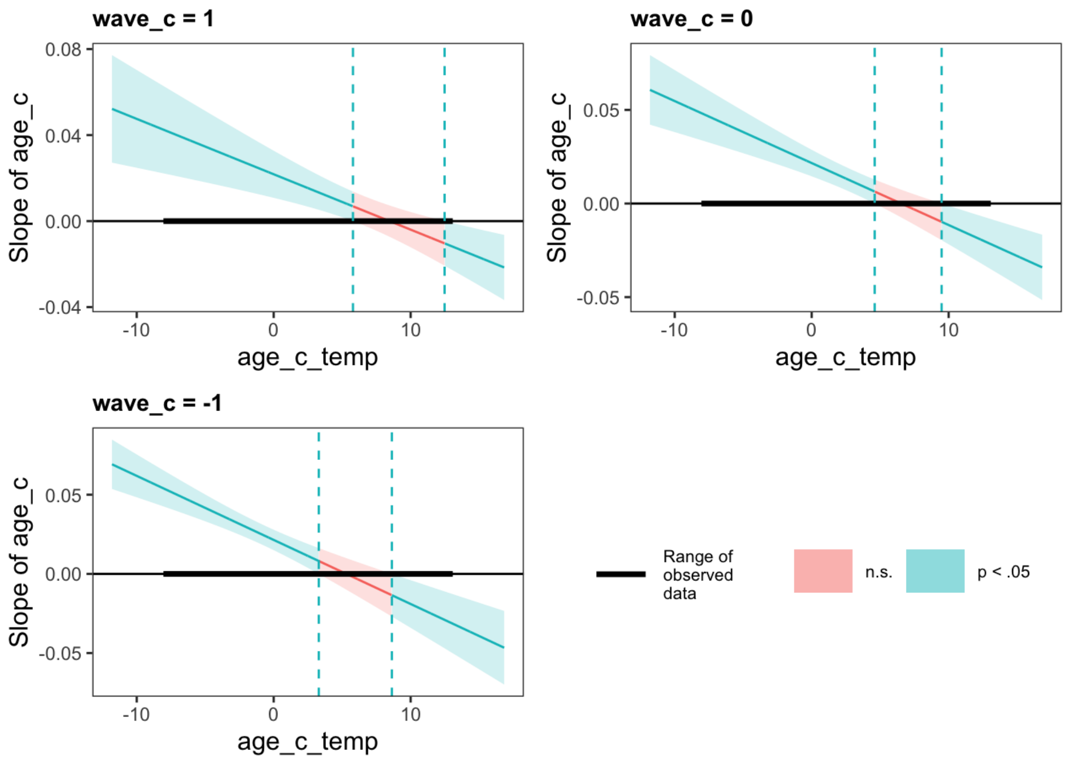


With standardized effects:

```
##                 term std.estimate  std.error      conf.low     conf.high
## 1                 fh  -0.00651459 0.02049034 -0.0466749095  0.0336457299
## 2                 sh   0.02136929 0.02048481 -0.0187801930  0.0615187701
## 3             wave_c   0.01085026 0.03050479 -0.0489380317  0.0706385549
## 4              age_c   0.16002004 0.04371478  0.0743406458  0.2456994359
## 5         I(age_c^2)  -0.08026442 0.04156155 -0.1617235572  0.0011947221
## 6          fh:wave_c  -0.01606514 0.02219917 -0.0595747118  0.0274444271
## 7          sh:wave_c  -0.04394964 0.02218244 -0.0874264184 -0.0004728692
## 8           fh:age_c  -0.04906794 0.02212475 -0.0924316545 -0.0057042276
## 9           sh:age_c   0.02532648 0.02212122 -0.0180303127  0.0686832733
## 10      wave_c:age_c  -0.03811260 0.02392096 -0.0849968146  0.0087716107
## 11     fh:I(age_c^2)   0.04674000 0.02366666  0.0003541946  0.0931257994
## 12     sh:I(age_c^2)  -0.02495371 0.02366972 -0.0713455073  0.0214380900
## 13 wave_c:I(age_c^2)   0.08269642 0.02074742  0.0420322332  0.1233606092
```

```
##                   term std.estimate  std.error    conf.low    conf.high
## 1                   fh  0.094032915 0.01739632  0.05993676  0.128129070
## 2                   sh  0.114787759 0.01739163  0.08070080  0.148874722
## 3               wave_c -0.044005399 0.02913437 -0.10110771  0.013096916
## 4                age_c  0.327175444 0.04923602  0.23067462  0.423676270
## 5           I(age_c^2) -0.260664374 0.04672174 -0.35223731 -0.169091442
## 6            fh:wave_c -0.035792062 0.01885622 -0.07274958  0.001165456
## 7            sh:wave_c -0.021964418 0.01884216 -0.05889437  0.014965537
## 8             fh:age_c  0.067585090 0.01877594  0.03078493  0.104385247
## 9             sh:age_c -0.004331794 0.01877294 -0.04112608  0.032462495
## 10        wave_c:age_c  0.001720267 0.02584893 -0.04894270  0.052383232
## 11       fh:I(age_c^2) -0.026795468 0.02008330 -0.06615801  0.012567077
## 12       sh:I(age_c^2) -0.002369001 0.02008589 -0.04173662  0.036998624
## 13 wave_c:I(age_c^2)  0.057057086 0.01783476  0.02210160  0.092012571
```

Which offer a significant improvement over the Main Effects-Only model:

```
## refitting model(s) with ML (instead of REML)
```

```
## Data: BTdat
## Models:
## main.effects.behav: learnrate ~ fh + sh + wave_c + age_c + I(age_c^2) + (1 | id)
## int.behav: learnrate ~ fh + sh + wave_c + age_c + I(age_c^2) + fh:wave_c +
## int.behav:     sh:wave_c + fh:age_c + sh:age_c + wave_c:age_c + fh:I(age_c^2) +
## int.behav:     sh:I(age_c^2) + wave_c:I(age_c^2) + (1 | id)
##                    Df     AIC     BIC logLik deviance  Chisq Chi Df Pr(>Chisq)
## main.effects.behav  8 -8078.7 -8026.9 4047.3  -8094.7
## int.behav          16 -8100.7 -7997.1 4066.4  -8132.7 38.029      8  7.437e-06
##
## main.effects.behav
## int.behav          ***
## ---
## Signif. codes:  0 '***' 0.001 '**' 0.01 '*' 0.05 '.' 0.1 ' ' 1
```

```
## refitting model(s) with ML (instead of REML)
```

```
## Data: BTdat
## Models:
## main.effects.brain: modrescale ~ fh + sh + wave_c + age_c + I(age_c^2) + (1 | id)
## int.brain: modrescale ~ fh + sh + wave_c + age_c + I(age_c^2) + fh:wave_c +
## int.brain:     sh:wave_c + fh:age_c + sh:age_c + wave_c:age_c + fh:I(age_c^2) +
## int.brain:     sh:I(age_c^2) + wave_c:I(age_c^2) + (1 | id)
##                    Df   AIC    BIC  logLik deviance  Chisq Chi Df Pr(>Chisq)
## main.effects.brain  8 74.49 126.30 -29.245    58.49
## int.brain          16 56.31 159.93 -12.155    24.31 34.179      8   3.77e-05
##
## main.effects.brain
## int.brain          ***
## ---
## Signif. codes:  0 '***' 0.001 '**' 0.01 '*' 0.05 '.' 0.1 ' ' 1
```

We can then plot the Interactions between Wave and the Quadratic Effect of Age:

And the Interactions between Age and the Effect Across Blocks in the First Half of the Task:

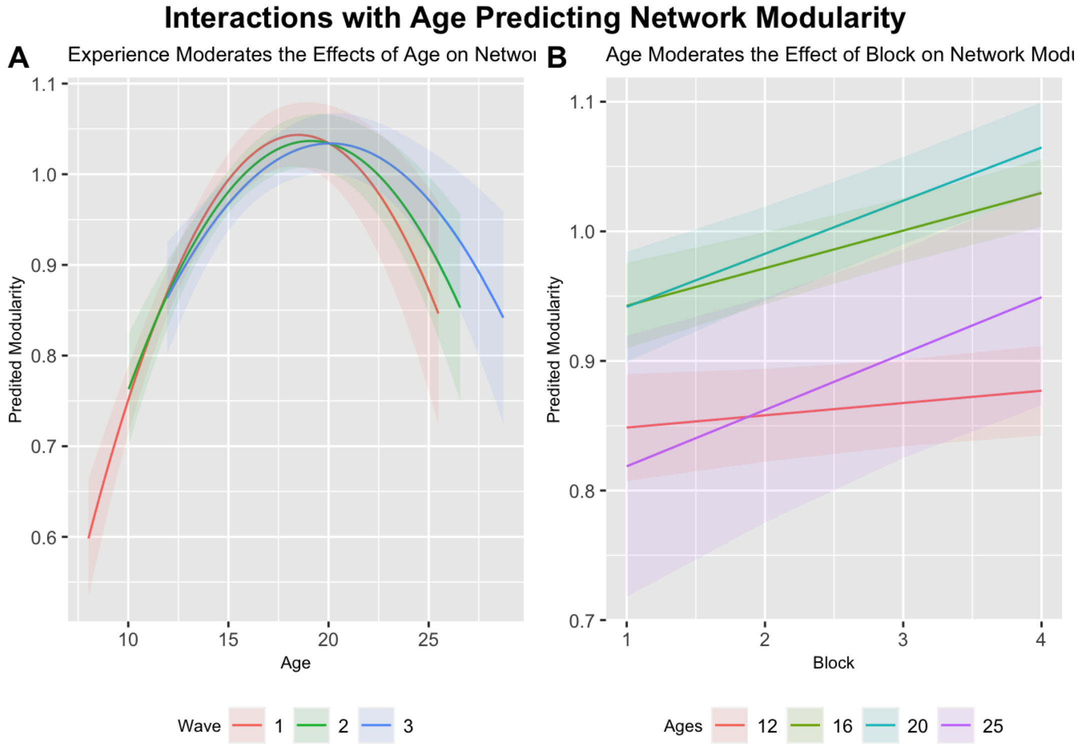


### Brain-as-Predictor Model

```
int.behav.brain = lmer(learnrate ~ fh + sh + wave_c + age_c + I(age_c^2) +
                        fh:wave_c + sh:wave_c + fh:age_c + sh:age_c + wave_c:age_c +
                        fh:I(age_c^2) + sh:I(age_c^2) + wave_c:I(age_c^2) +
                        modrescale_c + modrescale_c:fh + modrescale_c:sh + modrescale:wave_c +
                        modrescale_c:age_c + modrescale_c:I(age_c^2) +
                        modrescale_c:age_c:wave_c + modrescale_c:I(age_c^2):wave_c + (1 | id),
                      data = BTdat)
```

| | Learning Rate | | | |
|---|---|---|---|---|
| *Predictors* | *Estimates* | *SE* | *P-Value* | *df* |
| Intercept | 0.935 | 0.005 | **<0.001** | 794.679 |
| First Half | -0.000 | 0.002 | 0.886 | 4476.805 |
| Second Half | 0.001 | 0.002 | 0.775 | 4482.941 |
| Wave | -0.008 | 0.010 | 0.401 | 4670.922 |
| Age | 0.005 | 0.001 | **<0.001** | 718.911 |
| Age^2 | -0.000 | 0.000 | **0.046** | 679.282 |
| First Half * Wave | -0.002 | 0.003 | 0.415 | 4481.722 |
| Second Half * Wave | -0.004 | 0.002 | 0.072 | 4485.072 |
| First Half * Age | -0.001 | 0.000 | **0.018** | 4472.145 |
| Second Half * Age | 0.000 | 0.001 | 0.822 | 4502.070 |
| Wave * Age | -0.001 | 0.001 | 0.172 | 987.428 |
| First Half * Age^2 | 0.000 | 0.000 | 0.073 | 4469.835 |
| Second Half * Age^2 | -0.000 | 0.000 | 0.604 | 4483.426 |
| Wave * Age^2 | 0.000 | 0.000 | **0.001** | 4754.347 |

| | | | | |
|---|---|---|---|---|
| Modularity | -0.005 | 0.011 | 0.627 | 4764.879 |
| Modularity * First Half | 0.002 | 0.006 | 0.786 | 4532.995 |
| Modularity * Second Half | 0.012 | 0.006 | **0.044** | 4545.020 |
| Modularity * Wave | 0.009 | 0.009 | 0.317 | 4774.206 |
| Modularity * Age | 0.006 | 0.002 | **0.007** | 4222.697 |
| Modularity * Age^2 | -0.000 | 0.000 | 0.888 | 4389.408 |
| Modularity * Wave * Age | 0.004 | 0.002 | 0.093 | 4776.952 |
| Modularity * Wave * Age^2 | -0.001 | 0.000 | **0.041** | 4724.872 |
| **Random Effects** | | | | |
| $\sigma^2$ | 0.01 | | | |
| $\tau_{00\ id}$ | 0.00 | | | |
| ICC | 0.19 | | | |
| $N_{id}$ | 297 | | | |
| Observations | 4799 | | | |
| Marginal $R^2$ / Conditional $R^2$ | 0.071 / 0.243 | | | |

```
BTdat$modresids = scale(residuals(int.brain), scale=F, center=T)
int.behav.brain2 = lmer(learnrate ~ fh + sh + wave_c + age_c + I(age_c^2) +
                        fh:wave_c + sh:wave_c + fh:age_c + sh:age_c + wave_c:age_c +
                        fh:I(age_c^2) + sh:I(age_c^2) + wave_c:I(age_c^2) +
                        modresids + modresids:fh + modresids:sh + modresids:wave_c +
                        modresids:age_c + modresids:I(age_c^2) +
                        modresids:age_c:wave_c + modresids:I(age_c^2):wave_c + (1 | id),
                      data = BTdat)
```

Testing simple slopes:

```
## [redacted] While age_c_temp (2nd moderator) = -3.96 (- 1 SD) [redacted]
##
## JOHNSON-NEYMAN INTERVAL
##
## The Johnson-Neyman interval could not be found. Is the p value for your
## interaction term below the specified alpha?
##
## SIMPLE SLOPES ANALYSIS
##
## When age_c = -3.96 (- 1 SD):
##
##                              Est.   S.E.   t val.      p
## -------------------------- ------- ------ -------- ------
## Slope of modrescale_c        -0.02   0.01    -1.65   0.10
## Conditional intercept         0.90   0.01   164.77   0.00
##
## When age_c =  0.00 (Mean):
##
##                             Est.   S.E.   t val.      p
## -------------------------- ------ ------ -------- ------
## Slope of modrescale_c        0.00   0.01     0.18   0.86
## Conditional intercept        0.94   0.00   245.40   0.00
##
## When age_c =  3.96 (+ 1 SD):
##
##                             Est.   S.E.   t val.      p
## -------------------------- ------ ------ -------- ------
## Slope of modrescale_c        0.02   0.01     2.40   0.02
## Conditional intercept        0.95   0.00   195.06   0.00
##
## [redacted] While age_c_temp (2nd moderator) =  0.00 (Mean) [redacted]
```

```
##
## JOHNSON-NEYMAN INTERVAL
##
## When age_c is OUTSIDE the interval [-4.10, 4.23], the slope of modrescale_c
## is p < .05.
##
## Note: The range of observed values of age_c is [-7.84, 12.87]
##
## SIMPLE SLOPES ANALYSIS
##
## When age_c = -3.96 (- 1 SD):
##
##                               Est.   S.E.   t val.      p
## -------------------------- ------- ------ -------- ------
## Slope of modrescale_c        -0.02   0.01    -1.65   0.10
## Conditional intercept         0.90   0.01   164.77   0.00
##
## When age_c =  0.00 (Mean):
##
##                              Est.   S.E.   t val.      p
## -------------------------- ------ ------ -------- ------
## Slope of modrescale_c        0.00   0.01     0.18   0.86
## Conditional intercept        0.94   0.00   245.40   0.00
##
## When age_c =  3.96 (+ 1 SD):
##
##                              Est.   S.E.   t val.      p
## -------------------------- ------ ------ -------- ------
## Slope of modrescale_c        0.02   0.01     2.40   0.02
## Conditional intercept        0.95   0.00   195.06   0.00
##
## [redacted] While age_c_temp (2nd moderator) =  3.96 (+ 1 SD) [redacted]
##
## JOHNSON-NEYMAN INTERVAL
##
## When age_c is OUTSIDE the interval [-5.10, 2.67], the slope of modrescale_c
## is p < .05.
##
## Note: The range of observed values of age_c is [-7.84, 12.87]
##
## SIMPLE SLOPES ANALYSIS
##
## When age_c = -3.96 (- 1 SD):
##
##                               Est.   S.E.   t val.      p
## -------------------------- ------- ------ -------- ------
## Slope of modrescale_c        -0.02   0.01    -1.65   0.10
## Conditional intercept         0.90   0.01   164.77   0.00
##
## When age_c =  0.00 (Mean):
##
##                              Est.   S.E.   t val.      p
## -------------------------- ------ ------ -------- ------
## Slope of modrescale_c        0.00   0.01     0.18   0.86
## Conditional intercept        0.94   0.00   245.40   0.00
##
## When age_c =  3.96 (+ 1 SD):
##
##                              Est.   S.E.   t val.      p
## -------------------------- ------ ------ -------- ------
## Slope of modrescale_c        0.02   0.01     2.40   0.02
## Conditional intercept        0.95   0.00   195.06   0.00
```

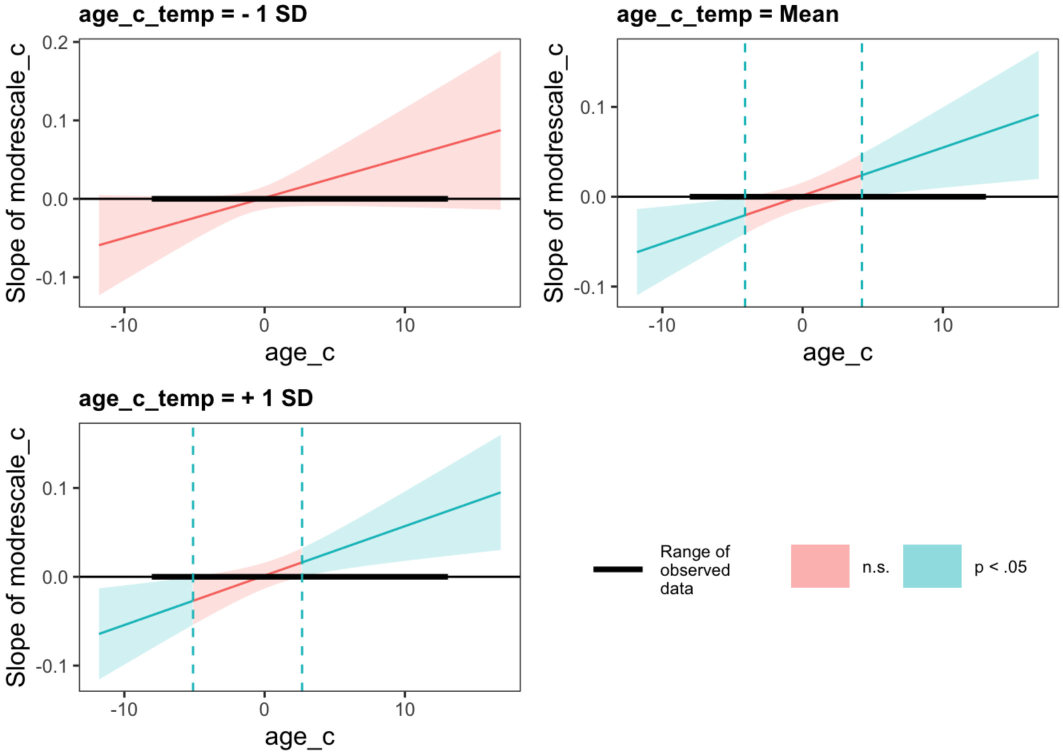


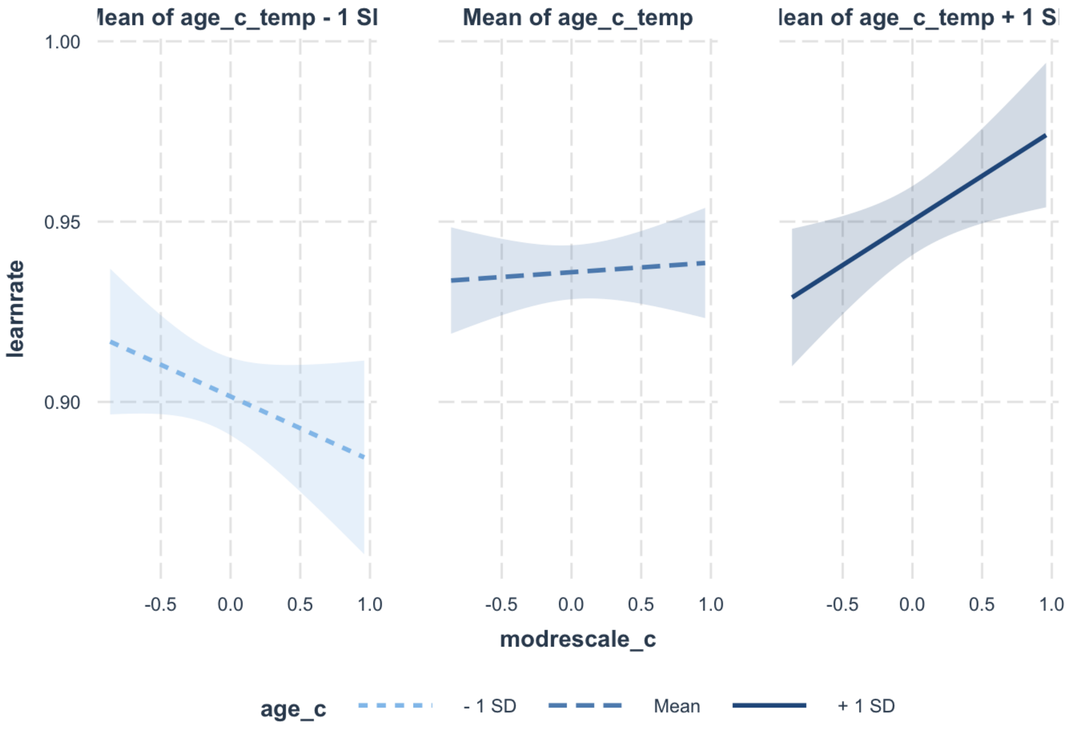


With standardized effects:

```
##                               term std.estimate  std.error     conf.low
## 1                               fh -0.002965629 0.02065077 -0.043440401
## 2                               sh  0.006120753 0.02145520 -0.035930672
## 3                           wave_c -0.056382133 0.06714436 -0.187982660
## 4                            age_c  0.174368375 0.04522131  0.085736233
## 5                       I(age_c^2) -0.085813312 0.04303595 -0.170162221
## 6                     modrescale_c -0.014178761 0.02914173 -0.071295495
## 7                        fh:wave_c -0.018222932 0.02236986 -0.062067052
## 8                        sh:wave_c -0.040122717 0.02228133 -0.083793324
## 9                         fh:age_c -0.054109015 0.02296495 -0.099119496
## 10                        sh:age_c  0.005252706 0.02335442 -0.040521121
## 11                    wave_c:age_c -0.033889061 0.02483320 -0.082561241
## 12                   fh:I(age_c^2)  0.044009260 0.02458309 -0.004172716
## 13                   sh:I(age_c^2) -0.012843476 0.02474955 -0.061351696
## 14               wave_c:I(age_c^2)  0.076376186 0.02248186  0.032312553
## 15                 fh:modrescale_c  0.005630874 0.02076889 -0.035075395
## 16                 sh:modrescale_c  0.044081878 0.02185564  0.001245604
## 17               wave_c:modrescale  0.061214707 0.06119269 -0.058720770
## 18              age_c:modrescale_c  0.062054717 0.02304264  0.016891970
## 19         I(age_c^2):modrescale_c -0.003551064 0.02520403 -0.052950049
## 20       wave_c:age_c:modrescale_c  0.035276261 0.02102099 -0.005924121
## 21 wave_c:I(age_c^2):modrescale_c -0.043386901 0.02125210 -0.085040252
##       conf.high
## 1   0.037509142
## 2   0.048172179
## 3   0.075218393
## 4   0.263000516
## 5  -0.001464403
## 6   0.042937973
## 7   0.025621189
## 8   0.003547889
## 9  -0.009098533
## 10  0.051026533
## 11  0.014783119
## 12  0.092191236
## 13  0.035664744
## 14  0.120439819
## 15  0.046337142
## 16  0.086918151
## 17  0.181150184
## 18  0.107217465
## 19  0.045847921
## 20  0.076476642
## 21 -0.001733549
```

Which offers a significant improvement over the Interaction Model:

```
## refitting model(s) with ML (instead of REML)
```

```
## Data: BTdat
## Models:
## int.behav: learnrate ~ fh + sh + wave_c + age_c + I(age_c^2) + fh:wave_c + 
## int.behav:     sh:wave_c + fh:age_c + sh:age_c + wave_c:age_c + fh:I(age_c^2) + 
## int.behav:     sh:I(age_c^2) + wave_c:I(age_c^2) + (1 | id)
## int.behav.brain: learnrate ~ fh + sh + wave_c + age_c + I(age_c^2) + fh:wave_c + 
## int.behav.brain:     sh:wave_c + fh:age_c + sh:age_c + wave_c:age_c + fh:I(age_c^2) + 
## int.behav.brain:     sh:I(age_c^2) + wave_c:I(age_c^2) + modrescale_c + modrescale_c:fh + 
## int.behav.brain:     modrescale_c:sh + modrescale:wave_c + modrescale_c:age_c + 
## int.behav.brain:     modrescale_c:I(age_c^2) + modrescale_c:age_c:wave_c + modrescale_c:I(age_c^2):wave_c + 
## int.behav.brain:     (1 | id)
##                 Df     AIC     BIC logLik deviance  Chisq Chi Df Pr(>Chisq)   
## int.behav       16 -8100.7 -7997.1 4066.4  -8132.7                            
## int.behav.brain 24 -8104.8 -7949.4 4076.4  -8152.8 20.101      8   0.009959 **
## ---
## Signif. codes:  0 '***' 0.001 '**' 0.01 '*' 0.05 '.' 0.1 ' ' 1
```

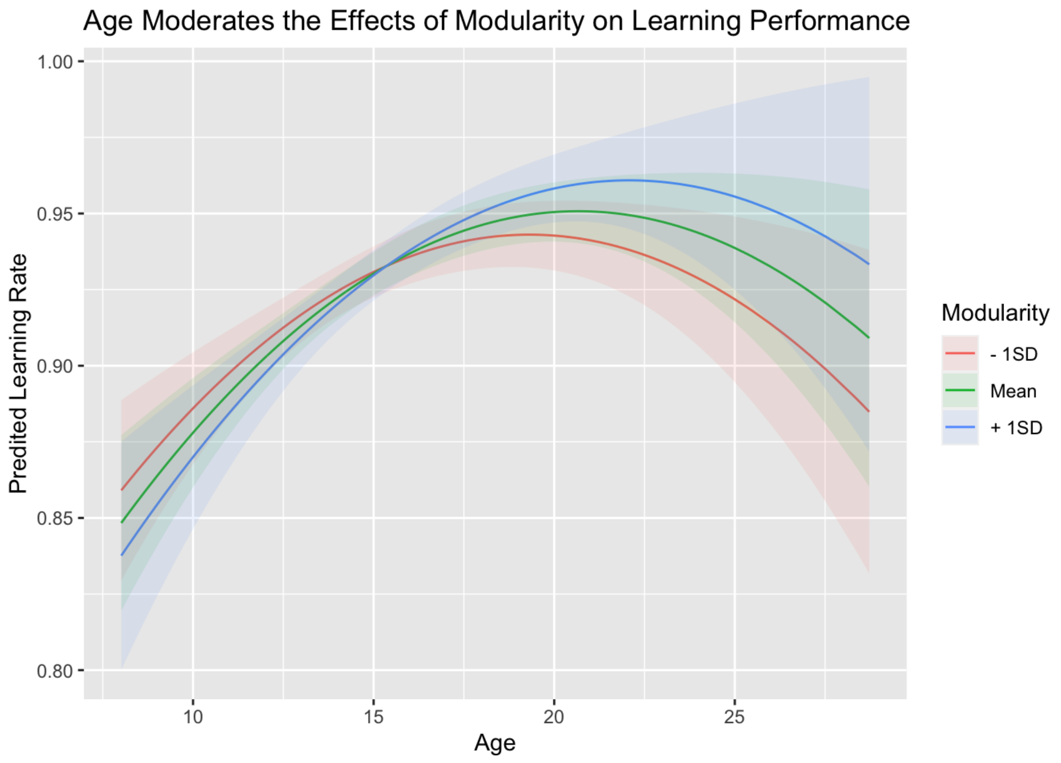


### Sensitivity Analyses:

Beta Regression

```
## Warning in nlminb(start = par, objective = fn, gradient = gr, control =
## control$optCtrl): NA/NaN function evaluation
```

```
##  Family: beta  ( logit )
## Formula:
## learnrate ~ fh + sh + wave_c + age_c + I(age_c^2) + fh:wave_c +
##     sh:wave_c + fh:age_c + sh:age_c + wave_c:age_c + fh:I(age_c^2) +
##     sh:I(age_c^2) + wave_c:I(age_c^2) + (1 | id)
## Data: BTdat_beta
##
##      AIC      BIC   logLik deviance df.resid
## -23560.4 -23456.7  11796.2 -23592.4     4783
##
## Random effects:
##
## Conditional model:
##  Groups Name        Variance Std.Dev.
##  id     (Intercept) 0.05362  0.2316
## Number of obs: 4799, groups:  id, 297
##
## Overdispersion parameter for beta family ():  4.5
##
## Conditional model:
##                     Estimate Std. Error z value Pr(>|z|)
## (Intercept)        2.6992672  0.0438614   61.54  < 2e-16 ***
## fh                 0.0019349  0.0224733    0.09  0.93139
## sh                -0.0170219  0.0226376   -0.75  0.45209
## wave_c             0.0174003  0.0414297    0.42  0.67449
## age_c              0.0261067  0.0098763    2.64  0.00821 **
## I(age_c^2)        -0.0032507  0.0016449   -1.98  0.04812 *
## fh:wave_c         -0.0008600  0.0253286   -0.03  0.97291
## sh:wave_c         -0.0336992  0.0257170   -1.31  0.19007
## fh:age_c          -0.0127162  0.0048629   -2.61  0.00892 **
## sh:age_c           0.0103354  0.0049578    2.08  0.03710 *
## wave_c:age_c      -0.0106581  0.0071246   -1.50  0.13466
## fh:I(age_c^2)      0.0013425  0.0008528    1.57  0.11546
## sh:I(age_c^2)     -0.0008065  0.0008574   -0.94  0.34693
## wave_c:I(age_c^2)  0.0031929  0.0009888    3.23  0.00124 **
## ---
## Signif. codes:  0 '***' 0.001 '**' 0.01 '*' 0.05 '.' 0.1 ' ' 1
```

```
## Warning in nlminb(start = par, objective = fn, gradient = gr, control =
## control$optCtrl): NA/NaN function evaluation
```

```
##  Family: beta  ( logit )
## Formula:
## learnrate ~ fh + sh + wave_c + age_c + I(age_c^2) + fh:wave_c +
##     sh:wave_c + fh:age_c + sh:age_c + wave_c:age_c + fh:I(age_c^2) +
##     sh:I(age_c^2) + wave_c:I(age_c^2) + modrescale_c + modrescale_c:fh +
##     modrescale_c:sh + modrescale:wave_c + modrescale_c:age_c +
##     modrescale_c:I(age_c^2) + modrescale_c:age_c:wave_c + modrescale_c:I(age_c^2):wave_c +
##     (1 | id)
## Data: BTdat_beta
##
##      AIC      BIC   logLik deviance df.resid
## -23556.0 -23400.5  11802.0 -23604.0     4775
##
## Random effects:
##
## Conditional model:
##  Groups Name        Variance Std.Dev.
##  id     (Intercept) 0.05265  0.2295
## Number of obs: 4799, groups:  id, 297
##
## Overdispersion parameter for beta family (): 4.51
##
## Conditional model:
##                                   Estimate Std. Error z value Pr(>|z|)
## (Intercept)                      2.6977772  0.0442695   60.94  < 2e-16 ***
## fh                               0.0059972  0.0226371    0.26  0.79107
## sh                              -0.0227138  0.0237548   -0.96  0.33898
## wave_c                          -0.0260457  0.0925483   -0.28  0.77838
## age_c                            0.0291126  0.0103666    2.81  0.00498 **
## I(age_c^2)                      -0.0033922  0.0017256   -1.97  0.04931 *
## modrescale_c                    -0.0300936  0.1078955   -0.28  0.78031
## fh:wave_c                       -0.0013795  0.0255528   -0.05  0.95695
## sh:wave_c                       -0.0321420  0.0258061   -1.25  0.21294
## fh:age_c                        -0.0149758  0.0050902   -2.94  0.00326 **
## sh:age_c                         0.0081099  0.0052703    1.54  0.12385
## wave_c:age_c                    -0.0099410  0.0075108   -1.32  0.18565
## fh:I(age_c^2)                    0.0014424  0.0008826    1.63  0.10220
## sh:I(age_c^2)                   -0.0007303  0.0008966   -0.81  0.41536
## wave_c:I(age_c^2)                0.0029811  0.0010733    2.78  0.00548 **
## fh:modrescale_c                  0.0673025  0.0639949    1.05  0.29294
## sh:modrescale_c                  0.0350278  0.0599459    0.58  0.55900
## wave_c:modrescale                0.0395954  0.0846701    0.47  0.64004
## age_c:modrescale_c               0.0480400  0.0200633    2.39  0.01665 *
## I(age_c^2):modrescale_c          0.0006118  0.0033190    0.18  0.85375
## wave_c:age_c:modrescale_c        0.0162742  0.0224119    0.73  0.46775
## wave_c:I(age_c^2):modrescale_c -0.0050642  0.0034373   -1.47  0.14067
## ---
## Signif. codes:  0 '***' 0.001 '**' 0.01 '*' 0.05 '.' 0.1 ' ' 1
```

Spline Models

```
gam.fit = gam(learnrate ~ s(wave_c, k=3) + s(age_c) + ti(age_c, wave_c, k=3, m=1) + s(id, bs="re"), data=BTdat)

gam.fit2 = gam(modrescale ~ s(wave_c, k=3) + s(age_c) + ti(age_c, wave_c, k=3, m=1) + s(id, bs="re"), data=BTdat)
```

```
## Summary:
##  * wave_c : numeric predictor; set to the value(s): -1.
##  * age_c : numeric predictor; with 30 values ranging from -7.843642 to 12.866360.
##  * id : factor; set to the value(s): BT0566. (Might be canceled as random effect, check below.)
##  * NOTE : The following random effects columns are canceled: s(id)
##
```

```
## Summary:
##  * wave_c : numeric predictor; set to the value(s): -1.
##  * age_c : numeric predictor; with 30 values ranging from -7.843642 to 10.096360.
##  * id : factor; set to the value(s): BT0566. (Might be canceled as random effect, check below.)
##  * NOTE : The following random effects columns are canceled: s(id)
##
```

```
## Summary:
##  * wave_c : numeric predictor; set to the value(s): 0.
##  * age_c : numeric predictor; with 30 values ranging from -5.833642 to 10.756360.
##  * id : factor; set to the value(s): BT0566. (Might be canceled as random effect, check below.)
##  * NOTE : The following random effects columns are canceled: s(id)
##
```

```
## Summary:
##  * wave_c : numeric predictor; set to the value(s): 1.
##  * age_c : numeric predictor; with 30 values ranging from -3.913642 to 12.866360.
##  * id : factor; set to the value(s): BT0566. (Might be canceled as random effect, check below.)
##  * NOTE : The following random effects columns are canceled: s(id)
##
```

```
## quartz_off_screen
##                 2
```

```
## Summary:
##  * wave_c : numeric predictor; set to the value(s): -1.
##  * age_c : numeric predictor; with 30 values ranging from -7.843642 to 12.866360.
##  * id : factor; set to the value(s): BT0566. (Might be canceled as random effect, check below.)
##  * NOTE : The following random effects columns are canceled: s(id)
##
```

```
## Summary:
##  * wave_c : numeric predictor; set to the value(s): -1.
##  * age_c : numeric predictor; with 30 values ranging from -7.843642 to 10.096360.
##  * id : factor; set to the value(s): BT0566. (Might be canceled as random effect, check below.)
##  * NOTE : The following random effects columns are canceled: s(id)
##
```

```
## Summary:
##  * wave_c : numeric predictor; set to the value(s): 0.
##  * age_c : numeric predictor; with 30 values ranging from -5.833642 to 10.756360.
##  * id : factor; set to the value(s): BT0566. (Might be canceled as random effect, check below.)
##  * NOTE : The following random effects columns are canceled: s(id)
##
```

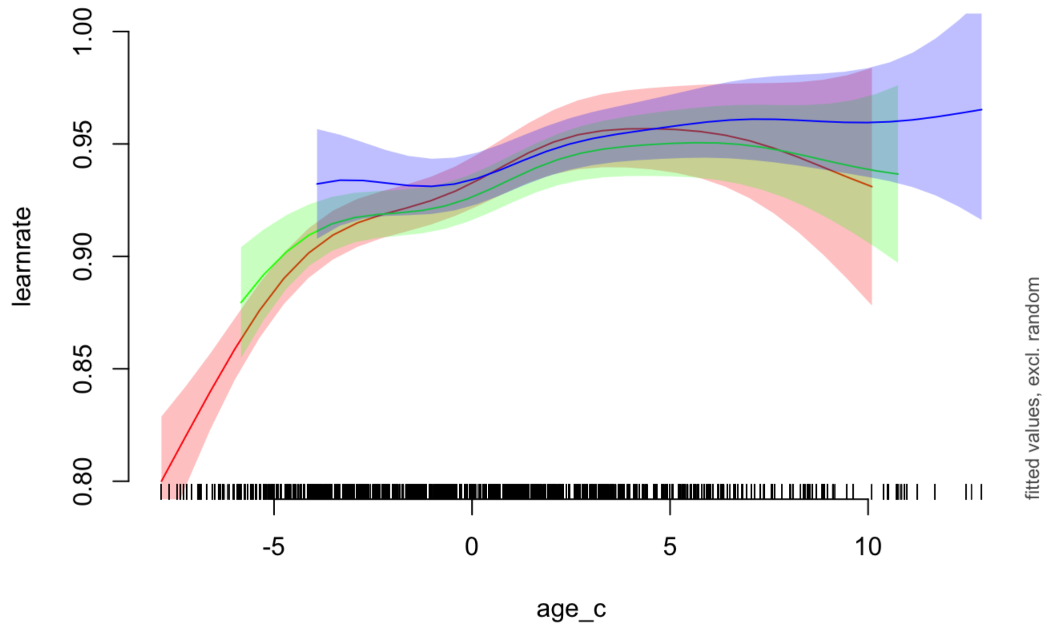

```
## Summary:
##  * wave_c : numeric predictor; set to the value(s): 1.
##  * age_c : numeric predictor; with 30 values ranging from -3.913642 to 12.866360.
##  * id : factor; set to the value(s): BT0566. (Might be canceled as random effect, check below.)
##  * NOTE : The following random effects columns are canceled: s(id)
##
```

```
## Summary:
##  * wave_c : numeric predictor; set to the value(s): -1.
##  * age_c : numeric predictor; with 30 values ranging from -7.843642 to 12.866360.
##  * id : factor; set to the value(s): BT0566. (Might be canceled as random effect, check below.)
##  * NOTE : The following random effects columns are canceled: s(id)
##
```

```
## Summary:
##  * wave_c : numeric predictor; set to the value(s): -1.
##  * age_c : numeric predictor; with 30 values ranging from -7.843642 to 10.096360.
##  * id : factor; set to the value(s): BT0566. (Might be canceled as random effect, check below.)
##  * NOTE : The following random effects columns are canceled: s(id)
##
```

```
## Summary:
##  * wave_c : numeric predictor; set to the value(s): 0.
##  * age_c : numeric predictor; with 30 values ranging from -5.833642 to 10.756360.
##  * id : factor; set to the value(s): BT0566. (Might be canceled as random effect, check below.)
##  * NOTE : The following random effects columns are canceled: s(id)
##
```

```
## Summary:
##  * wave_c : numeric predictor; set to the value(s): 1.
##  * age_c : numeric predictor; with 30 values ranging from -3.913642 to 12.866360.
##  * id : factor; set to the value(s): BT0566. (Might be canceled as random effect, check below.)
##  * NOTE : The following random effects columns are canceled: s(id)
##
```

```
## quartz_off_screen
##                 2
```

```
## Summary:
##  * wave_c : numeric predictor; set to the value(s): -1.
##  * age_c : numeric predictor; with 30 values ranging from -7.843642 to 12.866360.
##  * id : factor; set to the value(s): BT0566. (Might be canceled as random effect, check below.)
##  * NOTE : The following random effects columns are canceled: s(id)
##
```

```
## Summary:
##  * wave_c : numeric predictor; set to the value(s): -1.
##  * age_c : numeric predictor; with 30 values ranging from -7.843642 to 10.096360.
##  * id : factor; set to the value(s): BT0566. (Might be canceled as random effect, check below.)
##  * NOTE : The following random effects columns are canceled: s(id)
##
```

```
## Summary:
##  * wave_c : numeric predictor; set to the value(s): 0.
##  * age_c : numeric predictor; with 30 values ranging from -5.833642 to 10.756360.
##  * id : factor; set to the value(s): BT0566. (Might be canceled as random effect, check below.)
##  * NOTE : The following random effects columns are canceled: s(id)
##
```

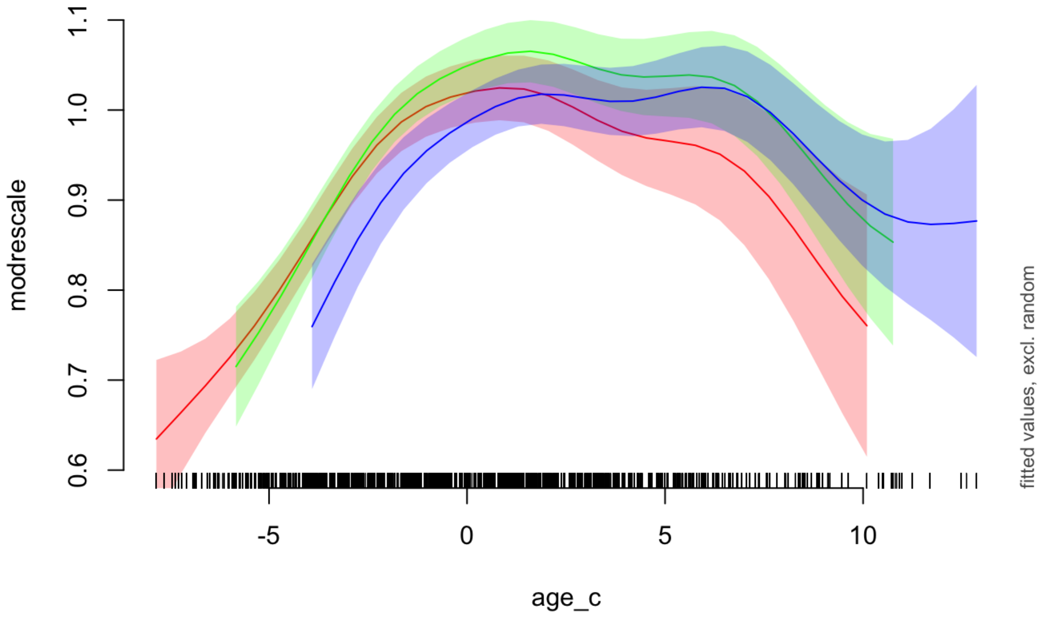


```
## Summary:
##  * wave_c : numeric predictor; set to the value(s): 1.
##  * age_c : numeric predictor; with 30 values ranging from -3.913642 to 12.866360.
##  * id : factor; set to the value(s): BT0566. (Might be canceled as random effect, check below.)
##  * NOTE : The following random effects columns are canceled: s(id)
##
```

Covariates: IQ + Gender

```
int.behav.covars = lmer(learnrate  ~ fh + sh + wave_c + age_c + I(age_c^2) +
                     fh:wave_c + sh:wave_c + fh:age_c + sh:age_c + wave_c:age_c +
                     fh:I(age_c^2) + sh:I(age_c^2) + wave_c:I(age_c^2) + iq + male + (1 | id),
                   data = BTdat)
summary(int.behav.covars)
```

```
## Linear mixed model fit by REML. t-tests use Satterthwaite's method [
## lmerModLmerTest]
## Formula: learnrate ~ fh + sh + wave_c + age_c + I(age_c^2) + fh:wave_c +
##     sh:wave_c + fh:age_c + sh:age_c + wave_c:age_c + fh:I(age_c^2) +
##     sh:I(age_c^2) + wave_c:I(age_c^2) + iq + male + (1 | id)
##    Data: BTdat
##
## REML criterion at convergence: -5685.8
##
## Scaled residuals:
##     Min      1Q  Median      3Q     Max
## -4.9198 -0.5132  0.2760  0.5969  3.2380
##
## Random effects:
##  Groups   Name        Variance Std.Dev.
##  id       (Intercept) 0.002676 0.05173
##  Residual             0.010230 0.10114
## Number of obs: 3602, groups:  id, 294
##
## Fixed effects:
##                     Estimate Std. Error         df t value Pr(>|t|)
## (Intercept)        8.140e-01  2.762e-02  1.325e+03  29.470  < 2e-16 ***
## fh                -8.583e-04  2.987e-03  3.273e+03  -0.287 0.773884
## sh                 2.090e-03  2.988e-03  3.273e+03   0.699 0.484307
## wave_c            -7.483e-03  7.645e-03  3.585e+03  -0.979 0.327754
## age_c              3.934e-03  1.492e-03  7.915e+02   2.636 0.008555 **
## I(age_c^2)        -2.171e-04  2.745e-04  9.220e+02  -0.791 0.429248
## iq                 1.114e-03  2.495e-04  1.360e+03   4.465 8.66e-06 ***
## male              -9.129e-03  6.982e-03  2.641e+02  -1.307 0.192206
## fh:wave_c         -1.492e-03  3.848e-03  3.273e+03  -0.388 0.698310
## sh:wave_c         -4.330e-03  3.850e-03  3.273e+03  -1.125 0.260814
## fh:age_c          -9.637e-04  5.155e-04  3.273e+03  -1.869 0.061662 .
## sh:age_c           6.014e-04  5.155e-04  3.273e+03   1.167 0.243485
## wave_c:age_c      -2.479e-03  1.262e-03  1.324e+03  -1.964 0.049749 *
## fh:I(age_c^2)      1.970e-04  9.983e-05  3.273e+03   1.974 0.048523 *
## sh:I(age_c^2)     -4.630e-05  9.985e-05  3.273e+03  -0.464 0.642920
## wave_c:I(age_c^2)  7.558e-04  2.066e-04  3.565e+03   3.658 0.000258 ***
## ---
## Signif. codes:  0 '***' 0.001 '**' 0.01 '*' 0.05 '.' 0.1 ' ' 1
```

```
##
## Correlation matrix not shown by default, as p = 16 > 12.
## Use print(x, correlation=TRUE)  or
##     vcov(x)        if you need it
```

```
int.iq = lmer(iq  ~ fh + sh + wave_c + age_c + I(age_c^2) +
                     fh:wave_c + sh:wave_c + fh:age_c + sh:age_c + wave_c:age_c +
                     fh:I(age_c^2) + sh:I(age_c^2) + wave_c:I(age_c^2) + male + (1 | id),
                   data = BTdat)
summary(int.iq)
```

```
## Linear mixed model fit by REML. t-tests use Satterthwaite's method [
## lmerModLmerTest]
## Formula: iq ~ fh + sh + wave_c + age_c + I(age_c^2) + fh:wave_c + sh:wave_c +
##     fh:age_c + sh:age_c + wave_c:age_c + fh:I(age_c^2) + sh:I(age_c^2) +
##     wave_c:I(age_c^2) + male + (1 | id)
##    Data: BTdat
##
## REML criterion at convergence: 23386.5
##
## Scaled residuals:
##     Min      1Q  Median      3Q     Max
## -3.5381 -0.5597 -0.0279  0.5733  3.6475
##
## Random effects:
##  Groups   Name        Variance Std.Dev.
##  id       (Intercept) 84.53    9.194
##  Residual             28.54    5.342
## Number of obs: 3602, groups:  id, 294
##
## Fixed effects:
##                     Estimate Std. Error         df t value Pr(>|t|)
## (Intercept)        1.066e+02  8.954e-01  3.503e+02 119.010  < 2e-16 ***
## fh                 1.455e-04  1.578e-01  3.294e+03   0.001 0.999264
## sh                 4.873e-04  1.578e-01  3.294e+03   0.003 0.997536
## wave_c            -1.632e+00  4.945e-01  1.692e+03  -3.301 0.000984 ***
## age_c             -1.073e-01  1.812e-01  3.506e+02  -0.593 0.553888
## I(age_c^2)         8.884e-02  3.195e-02  3.668e+02   2.780 0.005711 **
## male              -2.987e-01  1.091e+00  2.869e+02  -0.274 0.784536
## fh:wave_c         -6.140e-04  2.033e-01  3.294e+03  -0.003 0.997590
## sh:wave_c         -2.056e-03  2.033e-01  3.294e+03  -0.010 0.991934
## fh:age_c           6.869e-06  2.723e-02  3.294e+03   0.000 0.999799
## sh:age_c           2.300e-05  2.723e-02  3.294e+03   0.001 0.999326
## wave_c:age_c      -2.623e-02  1.332e-01  4.189e+02  -0.197 0.843955
## fh:I(age_c^2)     -1.116e-05  5.273e-03  3.294e+03  -0.002 0.998312
## sh:I(age_c^2)     -3.736e-05  5.274e-03  3.294e+03  -0.007 0.994348
## wave_c:I(age_c^2)  1.222e-02  1.127e-02  3.365e+03   1.084 0.278374
## ---
## Signif. codes:  0 '***' 0.001 '**' 0.01 '*' 0.05 '.' 0.1 ' ' 1
```

```
##
## Correlation matrix not shown by default, as p = 15 > 12.
## Use print(x, correlation=TRUE)  or
##     vcov(x)        if you need it
```

```
plot(ggeffect(int.iq,  terms=c('age_c [all]','wave_c [-1,0]')))
```

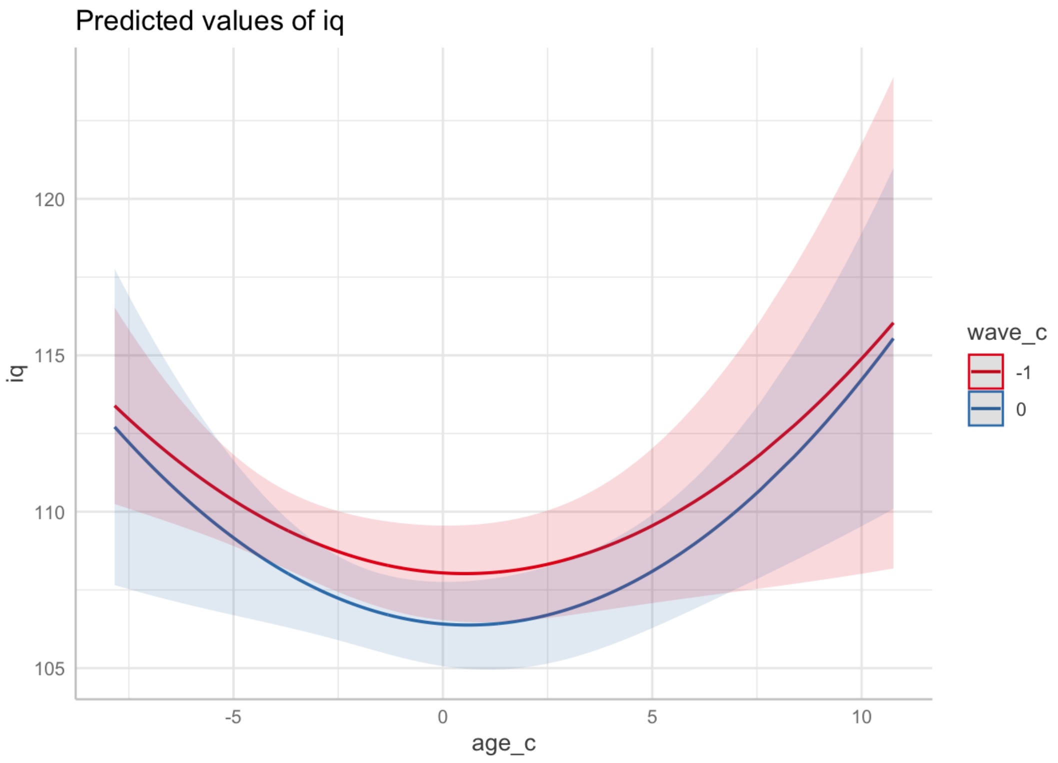


Puberty

```
cor(BTdat$age, BTdat$pds, use='complete.obs')
```

```
## [1] 0.8045119
```
